# Through the lens of causal inference: Decisions and pitfalls of covariate selection

**DOI:** 10.1101/2024.01.11.575211

**Authors:** Gang Chen, Zhengchen Cai, Paul A. Taylor

## Abstract

The critical importance of justifying the inclusion of covariates is a facet often overlooked in data analysis. While the incorporation of covariates typically follows informal guidelines, we argue for a comprehensive exploration of underlying principles to avoid significant statistical and interpretational challenges. Our focus is on addressing three common yet problematic practices: the indiscriminate lumping of covariates, the lack of rationale for covariate inclusion, and the oversight of potential issues in result reporting. These challenges, prevalent in neuroimaging models involving covariates such as reaction time, demographics, and morphometric measures, can introduce biases, including overestimation, underestimation, masking, sign flipping, or spurious effects.

Our exploration of causal inference principles underscores the pivotal role of domain knowledge in guiding co-variate selection, challenging the common reliance on statistical measures. This understanding carries implications for experimental design, model-building, and result interpretation. We draw connections between these insights and reproducibility concerns, specifically addressing the selection bias resulting from the widespread practice of strict thresholding, akin to the logical pitfall associated with “double dipping.” Recommendations for robust data analysis involving covariates encompass explicit research question statements, justified covariate inclusions/exclusions, centering quantitative variables for interpretability, appropriate reporting of effect estimates, and advocating a “highlight, don’t hide” approach in result reporting. These suggestions are intended to enhance the robustness, transparency, and reproducibility of covariate-driven analyses, encompassing investigations involving consortium datasets such as ABCD and UK Biobank. We discuss how researchers can use a transparent depiction of the covariate relationships to enhance the ethos of open science and promote research reproducibility.

## 1 Introduction

The pursuit of causality is a central objective in scientific inquiry, typically achieved through well-designed experiments or observational studies. Causal relationships manifest when one variable exerts an influence on another. Experiments aim to explore the impact of explanatory variables (exposures or treatments) on the response (or outcome) variable under diverse conditions. An alternative investigative approach is association studies, particularly valuable when discerning cause and effect is challenging. Association studies often serve as a preliminary step before establishing causal relationships. Both causal and association studies demand meticulous attention to covariate selection, a ubiquitous consideration in data analysis.

A causal effect denotes the influence that an explanatory variable exerts on the response variable. It signifies the alteration in the response that can be attributed to the manipulation or change in the explanatory variable, even when considering counterfactual scenarios, while keeping all other factors constant. In statistical analysis, the causal effect is quantified as the observed change in the response corresponding to variations in the explanatory variable. This analytical approach enables researchers to disentangle the specific contribution of the explanatory variable to changes in the response, while controlling for potential confounding factors. Understanding causal effects is fundamental to unraveling the underlying mechanisms of diverse phenomena and interventions.

In this context, a *covariate* is defined as a variable accompanying the explanatory variable in a statistical model. This explicit definition ensures consistency and clarity, avoiding the varied and sometimes offhand or pejorative terminologies such as control/adjustment variables, regressors/variables of no interest, nuisance regressors/variables, and extraneous covariates. Though often termed confounds in neuroimaging, it is crucial to note that confounds carry a distinct technical meaning, which we will explore further. Neuroimaging data analysis routinely involves covariates such as task conditions, participants, groups, estimated head motion, outliers, physiological recordings, age, sex, reaction time, global signal, and gray matter volume. Common practices, like excluding data points or participants due to outliers (e.g., excessive head motion), can be conceptually understood as a form of covariate integration. Sometimes, an explanatory variable assumes dual roles, serving as both a variable of interest and a covariate for another explanatory variable.

Neuroimaging studies exhibit various levels and degrees of variability, some meticulously controlled by the researcher, such as manipulated tasks in a factorial within-individual structure. However, uncontrollable incidents, including head motion and signals unrelated to neuronal activities, also influence the data. Between-individual variations, such as demographic factors (age, race, education, socioeconomic status), behavioral traits (reaction time), and physiological characteristics (head size), further contribute to the complexity. Faced with such diversity, the essential question for researchers is, *Which covariates should be included in the model to effectively address the study question at hand?*

### 1.1 Model considerations: a motivating example

The following presents a practical example from a neuroimaging study focusing on population-level modeling with data collected from participants spanning a wide age range. Assume that T1-weighted structural fMRI images are obtained when participants lay still in the scanner with no tasks involved. The behavior measure of short-term memory (STM) serves as the response variable. Additionally, six potential covariates are considered: voxel-level gray matter density (GMD) estimated with T1-weighted images, sex, age, intracranial volume (ICV), APOE genotype, and body weight. The primary objective of the study is to assess the causal relationship between GMD and STM. This scenario prompts several critical questions we address in this paper:

#### 1) Explanatory variable versus response variable

While it is biologically plausible to consider GMD as an explanatory variable and STM as the response variable, conventional neuroimaging tools often assume a response variable at the spatial-unit (e.g., voxel) level, with covariates that remain constant across spatial units. Is it justifiable, for the sake of tool convenience, to designate GMD as the response variable and STM as the explanatory variable?

#### 2) Covariates

When assessing the association between GMD and STM, should all six covariates be included in the model? Indiscriminately incorporating all available variables can lead to misleading inferences. How does one determine which covariates should or should not be incorporated? Can the importance of a covariate or model comparisons using step-up/down procedures guide its inclusion?

#### 3) Result interpretability/reporting

Typically, a single model (e.g., GLM) is used for a given dataset, producing an array of parameter estimates for all relevant covariates. However, is it appropriate to present all these estimates (e.g., sex differences in STM and the relationship between age and STM)?

#### 4) Experimental Design

In hindsight, could the experiment have been designed more efficiently? Which variables could have been omitted to prevent unnecessary data collection effort and resource wastage? Alternatively, what other variables might have been included to enhance effect estimation?

This example is to some extent an observational study. The explanatory variable GMD, being estimated solely from structural data, lacks the manipulability of variables seen in randomized controlled experiments. This raises a question about the feasibility of estimating the GMD-STM causal relationship. Moreover, if the focus shifts to relationship estimation akin to an association study, does this analytical pivot shield the investigator from, or render the investigator immune to, the various concerns and perspectives that inform how we approach the four aforementioned questions?

### 1.2 Covariate selection in models

Covariate selection is an essential process in statistical modeling, yet it is frequently underappreciated and prone to improper guidelines. For example, head size or brain volume is frequently used as a covariate. The question arises: is the adoption of such a quantity justified? In fact, the consideration of covariates often receives insufficient attention, with their impact sometimes regarded as a mere conventional practice or perhaps simply that “more is always better.” For instance, a survey on structural MRI analysis revealed a wide range in the number of covariates, varying from 0 to 14, with 37 unique ones employed across 68 models (Hyatt et al., 2020), and the reasoning behind collection and inclusion was seldom provided. This issue becomes particularly pronounced when a multitude of covariates, including demographic information, are readily available from large consortium datasets (e.g., ABCD, UK Biobank). This abundance can lead to the daunting choice among hundreds or thousands of covariates (Saragosa-Harris et al., 2022; Alfaro-Almagro et al., 2021; Zhao et al., 2023; Smith and Nichols, 2018), and readers should understand the potential consequences of such extreme cases.

The following are some common reasons that covariates are often included:

#### 1) Availability

A measure was acquired, and it might be related to the effects under study.

#### 2) Precedence

The variable has been widely used as a covariate previously, or within a particularly relevant prior study.

#### 3) Statistical evidence

The inclusion of a variable is driven by statistical metrics such as *p*-value, coefficient of determination *R*^2^, variance inflation factor, and information criteria. Additionally, a variable may be incorporated due to the extent of its association with other variables. In some cases, the percentage of variance explained by each of the considered covariates has been utilized and ranked to assess their “worthiness” in the employed model. Stepwise procedures are taught in some textbooks as a means of automating variable selection based on the statistical strength of each variable in a data-driven manner.

However, it is essential to critically evaluate such strategies. For instance, can the inclusion of a covariate be justified solely based on its statistical strength? Moreover, adding a covariate may either strengthen or weaken the statistical evidence for the effect of interest. By what principle could we differentiate what is more reasonable? These considerations give rise to three questions. First, do all covariates impact the effect of interest (or estimand) in the same way (and if not, how do we categorize their impacts)? Second, can a model be solely evaluated based on its own output? Third, how does the inclusion of a covariate affect interpretation of the model results? We advocate for the adoption of independent principles or criteria to determine whether a covariate should be included, in order to avoid circularity or interpretational issues. In fact, prior information and contextual knowledge are often available to inform the decision-making process. As we discuss further below, including a covariate without accounting for its underlying causal relations with the explanatory variable and response variables may lead to erroneous interpretations.

### 1.3 Modeling goals: prediction and inference with accuracy and precision

Prediction and inference represent two distinct yet crucial objectives in statistical modeling. *Prediction* employs a model to forecast future outcomes, emphasizing accuracy in predicting the response variable with less concern for the underlying relationships among input variables or features. Model comparison and variable selection are guided by the overall predictive performance of each model. In this context, the inclusion of numerous variables is feasible, and concerns about variable relationships hold less relevance. The emphasis in prediction is on achieving high predictive accuracy rather than uncovering causal relationships. Therefore, a model can be constructed to make predictions based on statistical correlations between variables, rather than establishing causal relationships. While correlated variables may influence each other, prediction does not necessitate an understanding of the underlying causal mechanisms. For instance, meteorologists predict weather conditions such as temperature, precipitation, and wind speed based on historical weather data, satellite images, and atmospheric variables. Their models focus on forecasting future weather patterns without necessarily understanding the underlying causal mechanisms driving weather phenomena.

On the contrary, *inference* aims to understand the relationships between an explanatory variable and the response variable. Consequently, all relationships among variables become important, and the essence of statistical inference lies in estimating the impact of a explanatory variable on the response variable. Natural questions arise: how to decide whether to include or exclude a covariate? How to characterize and leverage the relationships among variables? Do parameter estimates for all variables in a single model yield interpretable and meaningful insights?

Our primary focus here is on effect estimation in statistical inference, and the results are characterized by two properties: the *accuracy* and *precision* of the relationship between the explanatory variable and response. Accuracy gauges the extent of systemic biases, such as under- or overestimation or incorrect sign, while precision assesses the degree of uncertainty or vagueness, often represented by the standard error, arising from sampling, measurement errors, and other sources of variability. Each of these metrics plays a pivotal role in evaluating causal effects. However, precision characterizes the estimation uncertainty, and in some cases, it can be enhanced through large sample sizes and multiple studies. In contrast, systematic bias poses a more serious concern, as it can distort interpretation and causal inference.

### 1.4 Structure of this commentary

The ultimate objective of scientific inquiry is to comprehend the data-generating process behind the phenomenon being studied. While associative relationships can provide insights, a thorough understanding of causal relationships is essential when investigating the underlying mechanisms. Despite the well-known adage “correlation does not imply causation,” recent advances in causal inference emphasize the profound impact of these relationships on effect estimation (Pearl, 2009). Being attentive to causal relationships based on domain knowledge can deeply influence experimental design and model interpretation. Conversely, neglecting them can introduce bias, compromise statistical evidence reliability, or lead to misleading inferences. In this commentary, we advocate for explicit articulation of causal relationships using graphical representations, aiming to benefit the neuroimaging community.

The structure of this commentary is organized as follows, with a particular focus on didactic exploration and motivation of key considerations for causal inference. The first part presents case studies with possible avenues for the model building process. We commence with two toy examples to underscore key statistical considerations, realistic choices, and their implications. Subsequently, bluewe zoom out and introduce key technical terms and concepts in causal inference, using the case study quantities for reference. This includes discussing directed acyclic graphs (DAGs) and expound upon three fundamental categories of covariate relationships: confounder, mediator, and collider. Additionally, we explore auxiliary parent/child relationships of a covariate relative to the explanatory variable and response variable. Applying a causal inference lens, we employ the DAG framework to analyze the motivating example introduced in subsection 1.1. Lastly, we conclude with a summary and provide general recommendations for covariate selection.

## 2 Two toy examples

We present two toy examples to illustrate and gain insights into the influence of a covariate. Readers are encouraged to modify the values and explore different scenarios using the publicly available code.^1^ Here, we consistently refer to the relationship between a covariate *C* and the impact of the explanatory variable *X* on the response *Y* as “conditioning on *C*.” Various terms are used in the literature to describe this process, including “adjust for,” “control for,” “covariate out,” “regress out,” “fix,” “remove,” and “partial out.” For clarity and consistency, we use the term “condition on,” aligning with the concept of conditional probability. This implies that when conditioning the analysis on a covariate, we assume that the covariate takes one of its specific values.

### 2.1 Toy example I: investigating girl-tree height

Consider a scenario where a tree is planted when a girl is born, and both the girl’s and the tree’s heights are recorded annually for a span of 17 years. With this dataset, we aim to investigate two fundamental relationships: first, the association between tree height and girl height, and second, the growth rates exhibited by both the girl and the tree over the time period.

#### 2.1.1 Q1: What is the association between tree height and girl height?

We start with a model for the relationship between the girl height (girl_*t*_) and the tree height (tree_*t*_):

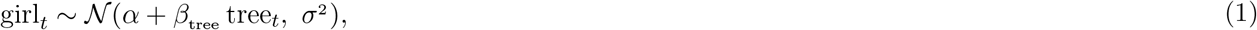

where α, *β*_tree_ and *σ* are the intercept, slope and standard deviation, respectively (*t* = 0, 1, …, 17 years). The model (1) can be more concisely expressed in an **R** compact symbolic formula girl ∼ tree. In the simulated data, we observe a strong linear correlation of height between tree and girl in Fig. 1, with a 95% quantile interval (0.099, 0.101) for *β*_tree_ and a coefficient of determination *R*^2^ ≈ 1.0 (Fig. 1B, red).

**Figure 1.**
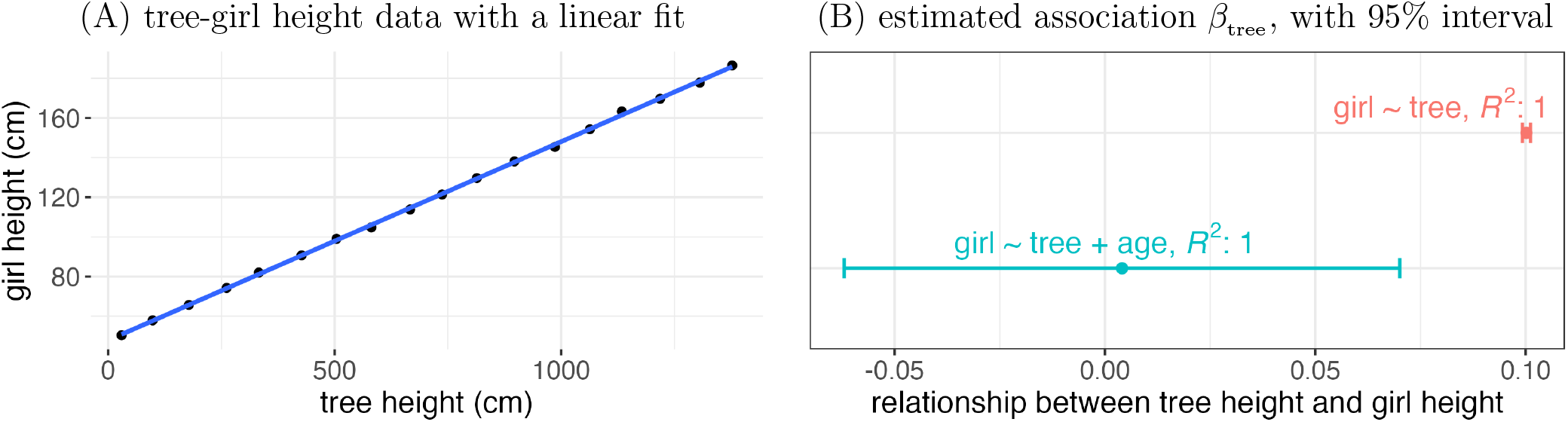
Toy example I - the association of growth between a girl and a tree. (A) Eighteen data points (black dots) were collected over 17 years for a girl and a tree. A linear relationship (blue line) can be seen between the two. (B) The simple regression (1), `girl ∼ tree`, indicates a growth ratio of one tenth with a 95% interval (0.099, 0.101) (red); conversely, the regression model (2), `girl ∼ tree + age`, indicates a near-zero ratio in the interval (−0.06, 0.07) (cyan).

Alternatively, we could feasibly construct another model `girl ∼ tree + age` with girl age included as a covariate:

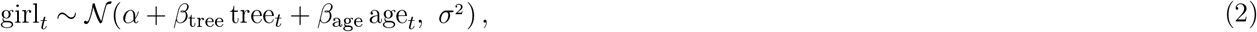

where age_*t*_ = *t*. In this case, the tree-girl relationship *β* tree appears to be essentially random noise with a 95% interval (−0.06, 0.07) (Fig. 1B, cyan). This model (2) also has a very high coefficient of determination *R*^2^ ≈ 1.0. When considering these two models, what determines which model is more appropriate for estimating an effect of interest? Can the strength of statistical evidence or model comparisons be used as justification for the inclusion of a covariate? If strength of statistical evidence is the decision-making criterion, then model (1) indicates that, when assessing the relationship between tree height and girl height, we should keep tree height while the model (2) suggests its removal. Model comparison would not help because the two models show very similar predictivity, even though the relations between the explanatory variable and the response variable are *very* different.

We note that both models are interpretable, but in different ways. Even though one may consider that the relationship revealed by the model (1) is correlative, not causal, the association is still informative: the girl’s height increases at about one-tenth the rate of the tree’s. In contrast, the model (2) shows that girl growth and tree growth are independent after conditioning on age. We can argue that both models are valid in their own way, but we might distinguish between them marginally based on practicality: at least in real life communication, information contained in the latter model (the girl’s growth is approximately 10 cm per year) is more tangibly conveyed than the former (the girl’s growth is about one-tenth that of the tree’s growth in her backyard).

#### 2.1.2 Q2: What is the growth rate of the girl?

Can the inclusion of a covariate be detrimental to effect estimation? Fig. 2A shows the girl’s height measurements versus age for the same data. To estimate the girl’s growth rate, we directly apply model (2) from above, yielding a 95% interval (2.5, 13.0) cm/year for *β*_age_ (Fig. 2B). This estimate exhibits remarkably poor precision and appears to be at odds with the visual assessment of the data in Fig. 2A.

**Figure 2.**
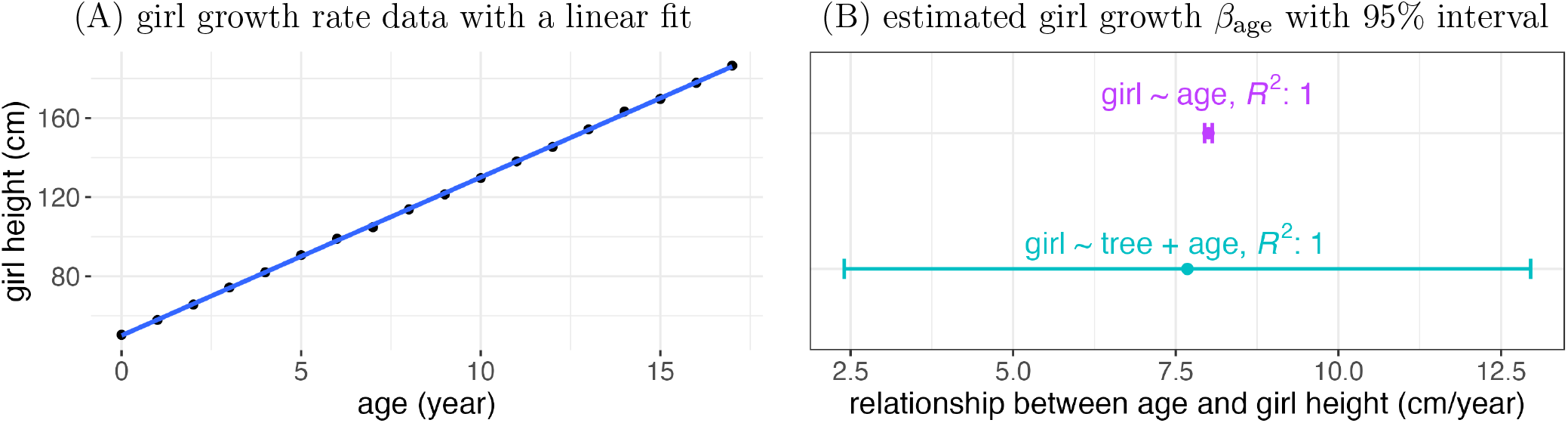
Girl growth rate, for same data as in Fig. 1. (A) Eighteen data points (black dots) were collected over 17 years for a girl and a tree. A virtually linear relationship (blue line) is shown for the girl growth rate. (B) A simple regression (3), `girl ∼ age`, indicates a 95% interval (7.95, 8.05) cm/year (purple) for the girl growth rate, compared to a much lower precision estimate (2.5, 13.0) (cyan) based on the regression model (2), `girl ∼ tree + age`.

We can also estimate the girl’s growth rate with a simpler regression by omitting the tree height as a covariate from the model (2), thus arriving at the model:

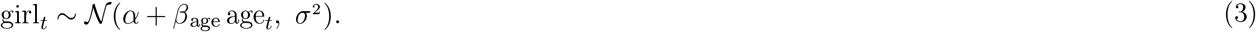

In this case, we observe a striking change in the estimated girl growth rate for the same data: a 95% interval (7.95, 8.05) cm/year (Fig. 2B, purple), again with *R*^2^ ≈ 1.0. Notably, this simpler regression (3) not only provides substantially improved estimation precision^2^ but also aligns more closely with the data visualization in Fig. 2A. Thus, the inclusion of a covariate may adversely affect estimation precision. We discuss this phenomenon and why it occurs, below.

#### 2.1.3 Three implications

This illustrative example unveils three pivotal insights. First, covariates wield nuanced impacts on estimation. In this context, the inclusion of age significantly enhances estimation accuracy, abolishing the previously observed strong relationship between tree height and girl height (Fig. 1). Conversely, the incorporation of tree height diminishes the precision of girl growth rate estimation (Fig. 2).

Second, the decision to include or exclude a variable transcends mere reliance on statistical metrics. Take models (1) and (2), for instance: even though their *R*^2^ values are indistinguishable, the interpretation of associated effects and covariate relationships can diverge. Each model holds unique utility, with the choice between them contingent on the specific research focus. While discerning these aspects and reasons for differing results is relatively straightforward in this toy case, real-world scenarios, especially in complex domains like neuroimaging, often present challenges in identifying variable relations. Underlying relationships may not be readily apparent, and some variables may be latent and not directly observable.

Third, relying on a single model might be inadequate for the proper estimation of all pertinent effects of interest. For instance, model (2), which incorporates age as a covariate, effectively removes the relationship between tree height and girl height but lacks precision when estimating girl growth rate compared to a straightforward regression (3) that excludes tree height as a covariate. This emphasizes the importance of considering distinct models to capture different effects of interest.

### 2.2 Toy example II: relationships among height, weight and sex

This example explores the relationships among height, weight, and sex in the adult population using a publicly available dataset (Davis, 1990) consisting of 199 adults, including 111 females (Fig. 3A). The dataset serves as a platform for us to investigate the intricacies of different modeling approaches when examining three effects of interest: the disparity in weight between sexes, the overall relationship between height and weight, and the difference in height between sexes.

**Figure 3.**
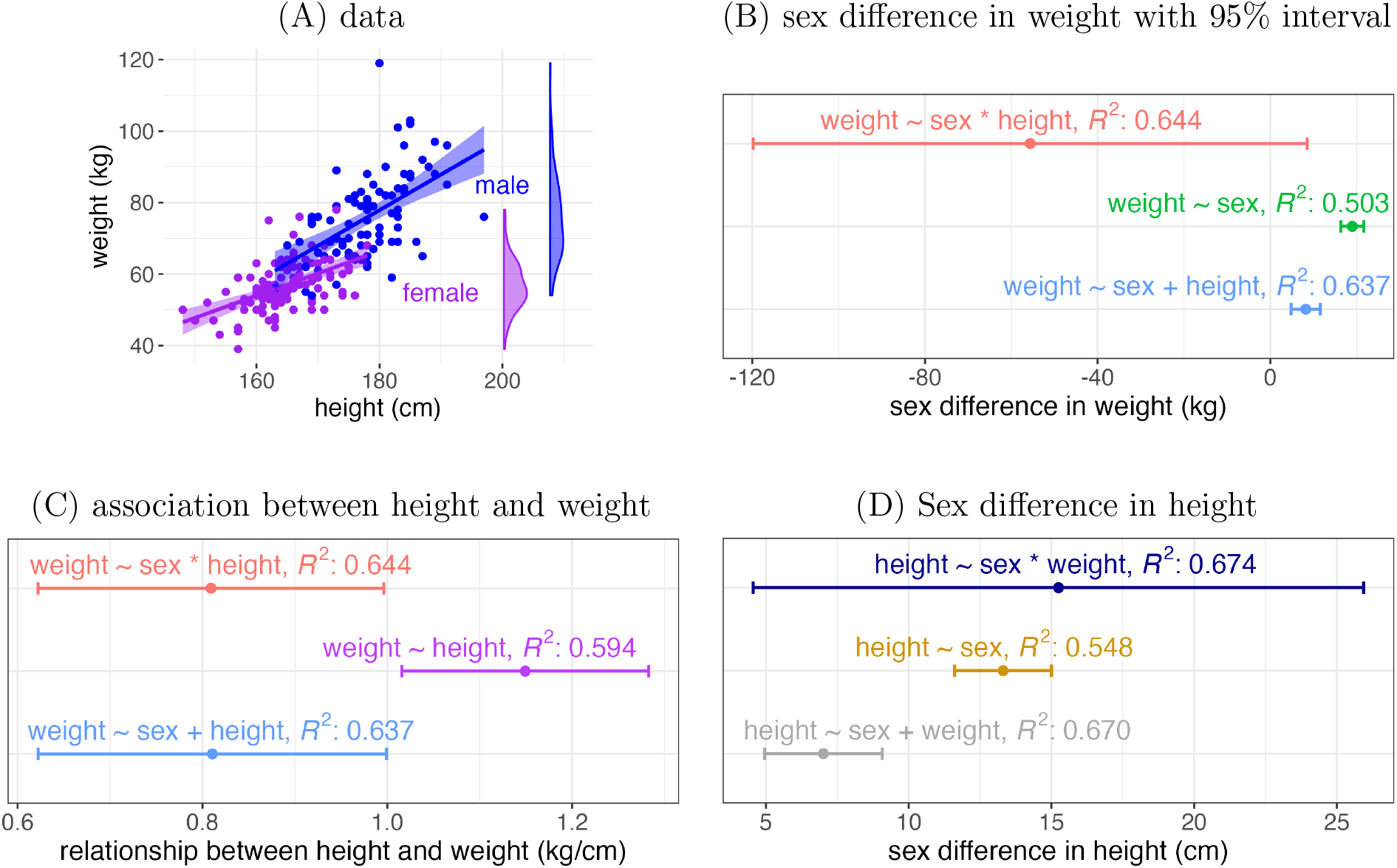
Sex, weight, and height relationships. (A) Real data from 199 participants are presented, with the shaded area indicating the 95% uncertainty band for the linear relationship within each sex. The density curves on the right-hand side illustrate that males (blue) generally have higher body weight than females (purple). (B-D) For the three effects of interest, point estimates are shown along with their corresponding 95% intervals for three distinct models.

#### 2.2.1 Q1: What is the sex difference in weight?

First, we simultaneously estimate the effects of sex and height on weight and also account for their interaction^3^,

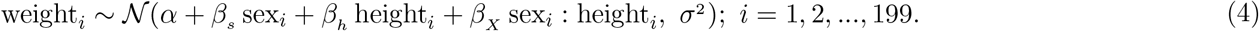

With sex_*i*_ dummy-coded as 0.5 for a male and −0.5 for a female, the weight difference *β*_*s*_ between sexes under the model (4) is estimated with a point estimate of −56 kg, a 95% interval (−120, 8) kg, and *R*^2^ = 0.644. Such a mostly negative estimate seems to contradict the common sense as well as the data visualization in Fig. 3.

As an alternative, we could use a straightforward model that corresponds to a conventional two-sample *t*-test,

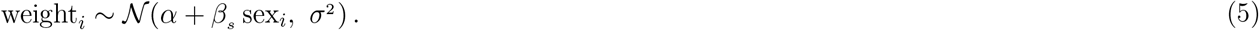

This yields a point estimate for *β*_*s*_ of 19.0 kg with a 95% interval (16.4, 22.6), which is more consistent with the data (Fig. 3B); the overall goodness of fit is lower than the previous model: *R*^2^ = 0.503.

Finally, we include height as a covariate but not the interaction between sex and height:

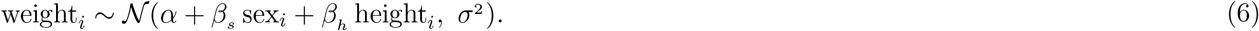

In this case, *β*_*s*_ has a point estimate of 8.2 kg, a 95% interval (4.8, 12.6), and *R*^2^ = 0.637 (Fig. 3B).

How can we evaluate each of these three models? The two-sample *t*-test (5) excludes height as a covariate and aligns with our intuitive expectations, although it produces the lowest coefficient of determination *R*^2^. While the most complex model (4) boasts the highest *R*^2^, its estimate contradicts common sense and the data. Model (6), which includes height as a covariate without an interaction, explains nearly the same proportion of data variability as model (4), but its estimate falls below that of the two-sample *t*-test (5). Common sense cannot help us differentiate the adequacy between the two, leaving a potentially inconclusive situation.

#### 2.2.2 Q2: What is the association between height and weight?

We adopt three distinct models to estimate the overall relationship between weight and height. Recycling the two models (4) and (6) leads to a virtually identical point estimate for *β*_*h*_ : 0.81 kg/cm, with a 95% interval (0.62, 1.00) kg/cm and *R*^2^ values provided above. However, opting for a straightforward linear regression,

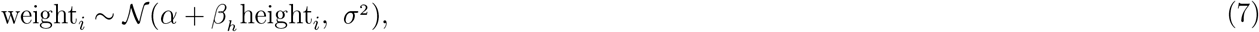

yields a different estimate for *β*_*h*_ : 1.15 kg/cm, with a 95% interval (1.02, 1.28) kg/cm and *R*^2^ = 0.594 (Fig. 3C). Again, some outcomes are similar and some are different, we do not have a straightforward means to critically differentiate between the three models (4), (6) and (7). As a consequence, the choice among these models remains unresolved. As displayed, perhaps the additional statistical evidence of the *R*^2^ would be the leading contender for guiding the decision-making process, but we have already seen issues with that criterion.

#### 2.2.3 Q3: What is the sex difference in height?

The estimation of the sex difference in height parallels the process applied to the sex difference in weight, as detailed in Section 2.2.1. We explore three distinct models, as follows:

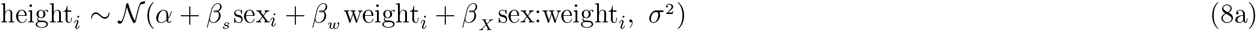

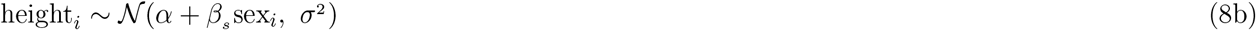

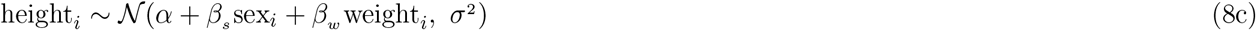

These models produce distinct outcomes (Fig. 3D). Their respective estimates for *βs* are 15.25, 13.31, and 7.01 cm, with respectively accompanying 95% intervals (4.55, 25.94), (11.61, 15.01), and (4.95, 9.07), and corresponding *R*^2^ values 0.674, 0.548, and 0.670. Selecting the most appropriate model remains unclear.

#### 2.2.4 Three implications

The three lessons we learned from the tree-girl example in Section 2.1 also apply to this toy example. First, the inclusion of a covariate can be detrimental and result in biased estimation (Fig. 3B,D). Second, statistical evidence based on a model alone cannot be used to determine its adequacy. In both sex difference in weight and height, the most complex models, (4) and (8a), have the highest *R*^2^, but the corresponding effect estimation is quite different and less precise than alternative models. Third, one cannot rely on a single complex model to estimate all relevant effects. Instead, separate models may be required for individual effects, as shown in the two-sample *t*-test models of (5) and (8b).

A fundamental question remains even in these simple toy models: how can we effectively evaluate the adequacy of a model? These examples vividly illustrate that conventional statistical metrics alone are inadequate for this task. Simply comparing models based on their ability to explain variability falls short of guaranteeing a meaningful answer. Instead, we need a principled way to determine when it is beneficial or detrimental to include a covariate. In the next section, we will discover that a more robust approach to differentiating models relies on external information. Specifically, it involves leveraging prior knowledge about the relationships among the explanatory variable, the response variable, and possible covariates.

## 3 Causal inference and directed acyclic graphs (DAGs)

One fundamental aspect of covariate selection relies on a comprehensive grasp of causal relationships, a concept extensively explored through causal inference (Pearl, 2009). This understanding provides a logical and consistent framework for addressing questions related to model structures, covariates, and interpretation. In this context, we apply causal inference to the two toy examples, demonstrating how it resolves issues and apparent discrepancies. Moreover, we posit that causal inference offers a valuable framework to assist researchers in experimental design and analyses, thereby enhancing study interpretability and reproducibility.

Unlike mere associations, causal relationships inherently possess directionality, indicating that influence flows from the cause to the effect, not vice versa. The directed acyclic graph (DAG) representation is central to illustrating the relationship between an explanatory variable *X* and a response *Y*. Thus, when establishing a causal connection between *X* and *Y*, it is visually represented as *X* → *Y*, signifying that *X* exerts influence on *Y*. In dealing with multiple variables, a DAG, like a flowchart-style structure (Greenland and Pearl, 2017), can be employed. This graphical tool serves to depict prior knowledge or hypothesized relationships among variables, offering an intuitive yet rigorously precise means to assess and present model constructions with conceptual clarity. DAGs can then be heuristically translated into model formulations, serving a similar graphical and organizational purpose to Feynman Diagrams in particle physics.

Establishing a causal relationship in observational studies, without the benefit of randomized control, presents challenges, yet remains feasible. DAGs serve as pivotal aids in this endeavor, visually mapping hypothesized causal relationships among variables. They furnish a structured framework for discerning directionalities and uncovering potential biases or alternative explanations. DAGs serve multiple purposes as invaluable tools: they elucidate hypotheses, pinpoint covariates for inclusion or exclusion, guide model construction, and facilitate sensitivity analysis. By offering a systematic and transparent approach to causal inference in observational studies, DAGs empower researchers to fortify the evidence supporting causal relationships among variables.

While the concept of causality has been a subject of debate and controversy throughout history, making it difficult to provide a precise definition, it remains fundamental to science and daily life. The exploration of causation can be traced back to ancient philosophers such as Democritus and Aristotle. However, rather than focusing on metaphysics (e.g., does time truly cause a girl to grow taller?), we adopt a practical approach aligned with the broader community of causal inference, embracing an empirical perspective. In this context, a causal relationship refers to a directional influence that the analyst assumes exists based on current knowledge.

### 3.1 Basic DAG terminology: causal relationships, path types and more

We note some of the most widely concepts in DAGs.

#### Components

Three components are associated with a DAG: *nodes, arrows*, and *paths*. Nodes (or vertices) represent the variables under investigation, while arrows (or arcs) symbolize the directional causal connections between these variables. By convention, the arrows are typically arranged in a consistent visual direction (e.g., left to right). A path is formed by a sequence of arrows, regardless of their direction, connecting the explanatory variable to the response variable (e.g., *X* ← *C* → *Y*, Fig. 4A).

**Figure 4.**
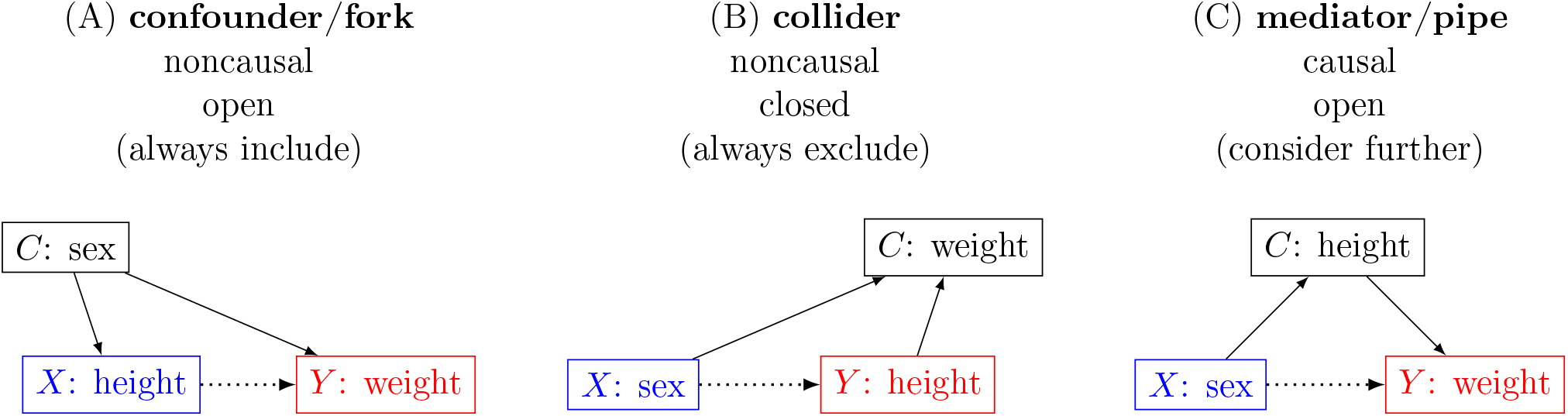
Three basic directed acyclic graph (DAG) types. The relationship under study is between an explanatory variable *X* (blue) and a response variable *Y* (red), as indicated by the dotted arrow. An arrow with a solid line indicates the *a priori* influence direction. A covariate *C* can be a confounder (A), a collider (B), or a mediator (C). The three variables of sex, height, and weight in toy example II are used to illustrate the three DAG types.

#### Relationships

There are three principal categories of covariate relationships, which are demonstrated in Fig. 4. The covariate *C* is a:

1. *confounder* (or fork) if it influences both *X* and *Y* (Fig. 4A) as a common cause (or common parent).
2. *collider* if it is influenced by both *X* and *Y* (Fig. 4B); that is, it is a common effect (or common child).
3. *mediator* (or pipe) if it is influenced by *X* while it also influences *Y* (Fig. 4C).

These three types provide a transparent way to convey the analyst’s prior information, theoretical constructs, and assumptions. The impact of each on the interpretation for the relationship between *X* and *Y* will be discussed in the next three subsections.

#### Path categories

Causal inference relies on the categorization of paths between *X* and *Y*. This can be done in two distinct ways, by distinguishing: between causal and noncausal paths, or between open and closed paths. A path is called *causal* when all arrows point away from *X* and towards *Y* (e.g., *X* → *C* → *Y*, as in Fig. 4C); otherwise, it is considered *noncausal* (e.g., *X* ← *C* → *Y* in Fig. 4A; *X* → *C* ← *Y* in Fig. 4B). Furthermore, a causal path can be either *direct* (i.e., *X* → *Y*) or *indirect* (e.g., *X* → *C* → *Y*, Fig. 4C). Paths within a DAG are categorized as open or closed depending on their impact on the response variable. An *open* path permits the response variable to vary freely, operating independently of other variables in the DAG. It lacks colliders and signifies potential causal relationships. Conversely, a *closed* path restricts the response variable from varying independently; it includes one or more colliders. A *backdoor* path is a non-causal path or one with an arrow entering the explanatory variable, while a *front-door* path is characterized by all arrows pointing away from the explanatory variable. A *minimally sufficient set* refers to the smallest set of variables that, when conditioned upon, effectively closes all open non-causal paths.

#### Parent/child relations

Four auxiliary cases are worth discussing. When a covariate *C* is connected with either the explanatory variable *X* or the response *Y, C* is a child (or descendant, or proxy; Fig. 5A-B) or a parent (or ancestor; Fig. 5C-D). In each of these four cases, *C* is referred to as *neutral* in the sense that it is not on a causal path between *X* and *Y*. Subtleties abound among these four cases. As a child of *X* (Fig. 5A), *C* is sometimes referred to as a post-treatment variable, and it is generally not recommended to use as a covariate because it typically leads to reduced precision (Montgomery et al., 2018; Gelman et al., 2020; McElreath, 2020) or to biases, especially when the explanatory variable contains measurement errors (Wysocki et al., 2022). When *C* is a child of *Y* (Fig. 5B), conditioning on *C* would incur biases (Cinelli et al., 2022). When *C* is a parent of *X* (Fig. 5C), it is a precision parasite because conditioning on *C* may lead to degraded precision. In contrast, *C* as a parent of *Y* (Fig. 5D) may help improve precision (Cinelli et al., 2022). For this reason, it is termed as a precision variable (Witte et al., 2020), additive exogenous covariate, or competing exposure.

**Figure 5.**
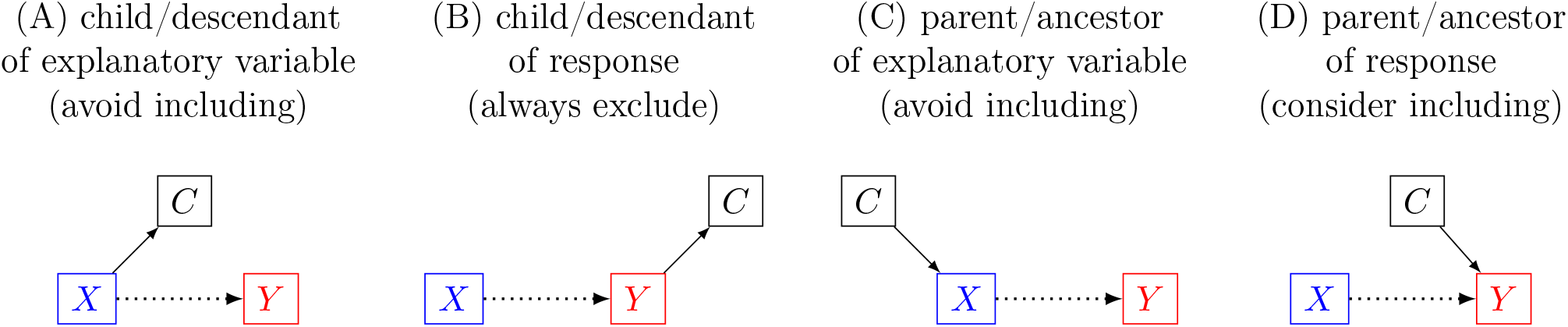
Covariate as a child or parent. The relationship under study is between an explanatory variable *X* (blue) and a response variable *Y* (red), as indicated by the dotted arrow. The explanatory variable *X* (A) or the response *Y* (B) may single-handedly influence the covariate *C*. Alternately, *C* may play the role of a parent, influencing *X* (C) or *Y* (D), but not both.

#### Causal effects

The value of a DAG rests in its direct representation of the underlying processes that generate the data. Causal relationships are characterized by paths via arrows that are always in the direction from the explanatory variable to the response variable (e.g., a path though a mediator *X* → *C* → *Y* in Fig. 4C). The *total causal effect* encompasses the combined influence over all causal paths. A *direct causal effect* signifies an impact that does not flow through any mediator.

#### Remarks

The decision to include a covariate will depend on its causal relationship with other variables, and these can be conveniently depicted within a DAG. This information resides “outside” of the model, and may involve discussions, debate and background information. Consider how in a linear regression, all variables are treated equally and their potentially complex interrelations are hidden, except perhaps by the *a posteriori* assessment of collinearity and amount of data variability–and even these ignore any notion of causality. In contrast, the prior knowledge exemplified in a DAG encapsulates causal relationships that can prove indispensable in untangling such complexities.

The relationships represented by a DAG possess a degree of abstraction. In essence, the directional influence denoted by an arrow does not convey an absolute meaning (Greenland and Pearl, 2017), but more of a notional one among the present variables. To illustrate, consider the causal link between a task condition (e.g., incongruent) and the resulting BOLD response, often perceived as a direct effect in routine data analysis. However, this “direct” effect is context-dependent; in general, we would omit intermediary variables such as cerebrovascular reactivity and neural responses, which act as mediators, for the sake of practicality.

### 3.2 Variable type: confounder

As a general rule, a confounder should always be conditioned on. A confounder *C* serves as a common cause for both *X* and *Y*. As such, apart from the direct causal path *X* → *Y*, the presence of the confounder *C* means there is also a noncausal, open path *X* ← *C* → *Y*. If *C* is not conditioned on, this noncausal path is not closed, and when interpreting the model results we cannot distinguish whether any observed association is a result of the direct causal path or of the influence of the noncausal path. This ambiguity will harm the accuracy and precision of effect estimation. Thus, it is essential to close the noncausal path by conditioning on the confounder, enabling one to adequately establish the causal effect of *X* on *Y*.

Understanding the role of a confounder is crucial for elucidating key aspects of model construction. Take, for instance, the classic phenomenon of *spurious correlation in ratios* (Pearson, 1896) in compositional data analysis. When three variables–*X, Y*, and *C*–are mutually independent, the correlation between *X* and *Y* is essentially zero. However, the correlation between the ratios *X/C* and *Y/C* can be remarkably high (e.g., 0.5 in cases with uniform distributions for each variable). What might seem like “spurious” correlation in the latter scenario arises due to the confounding role of *C*. Another popular instance of overlooking a confounder is illustrated by Simpson’s paradox (Pearl and Mackenzie, 2018). In general, biased estimation due to the failure of conditioning on a confounder can occur in various directions such as overestimation, underestimation, masking a true effect, sign reversal, and spurious effect (Appendix A.1).

#### 3.2.1 Revisiting toy example I

Model comparisons within each question in toy example I in Sec. 2.1 demonstrate the impact of a confounder. In Q1, when tree height is the explanatory variable *X* and girl height acts as the response *Y*, age is a confounder (Fig. 6A). As such, it is vital to include age in the model (2), `girl ∼ tree + age`, when one examines whether a *causal effect* exists between tree and girl heights (Fig. 6C). The exclusion of age, as shown in the model (1), `girl ∼ tree`, would only reveal the noncausal effect of the tree height on girl height (Fig. 6C). In essence, tree height is predictive of girl height, but it does not establish a causal relationship.

**Figure 6.**
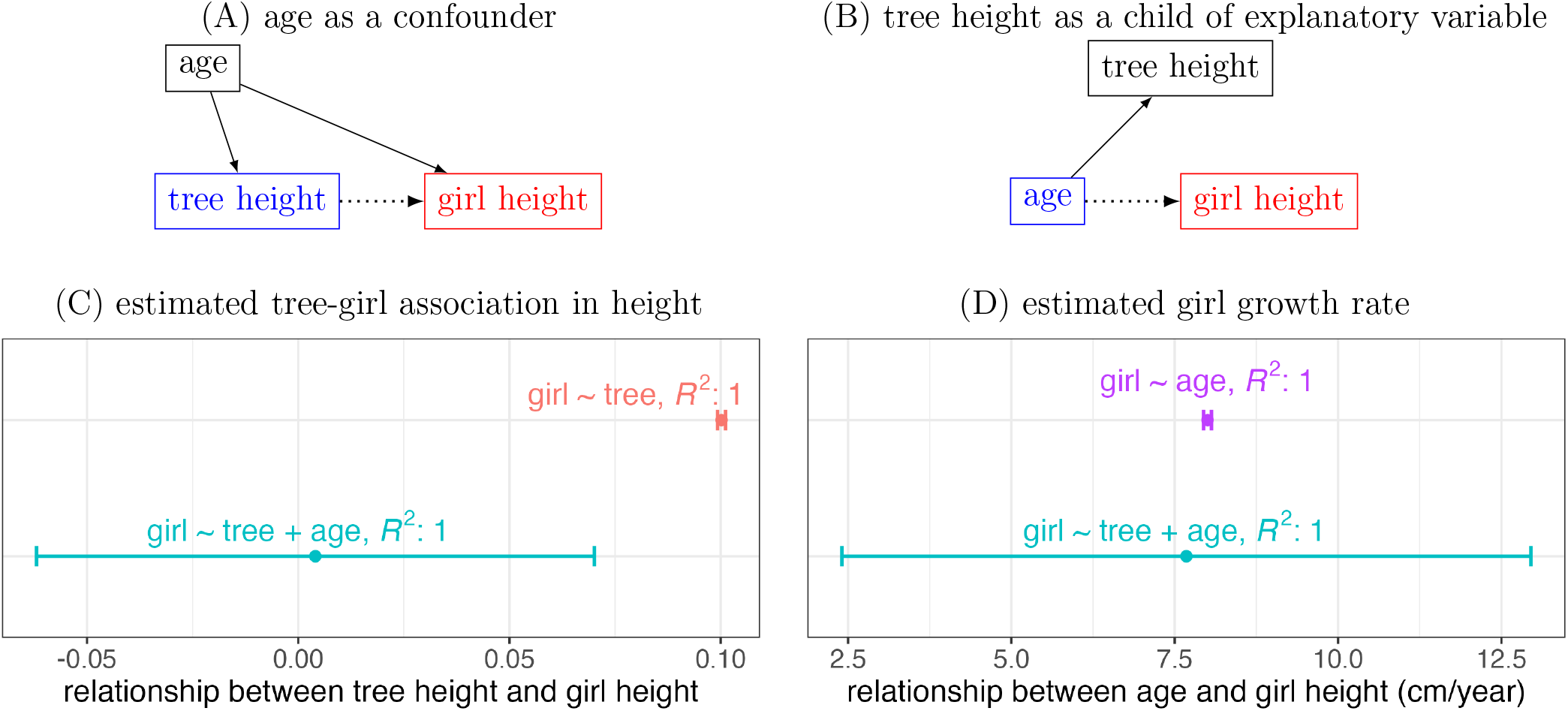
Toy example I revisited. In the top row, the explanatory variable and the response variables are indicated in blue and red, respectively. The relationship under study is indicated with a dotted-line arrow. (A) Age acts as a confounder between tree height and girl height. Thus, it is important to condition on age in assessing the causal relationship. (B) Tree height becomes a descendant of the explanatory variable when the focus is on girl growth rate. Such a variable is generally not recommended as a covariate. (C) and (D) are estimated results, the same as Figs. 1B and 2B.

In Q2 of toy example I, we see how estimation precision can be harmed if the descendant of the explanatory variable is included as a covariate. When the effect of interest changes to the girl growth rate, tree height becomes a child of the explanatory variable (Fig. 6B). Therefore, conditioning on tree height, as in the model (2), `girl ∼ tree + age`, leads to an estimation with poor precision (Fig. 6D) compared to the simple regression (3), `girl ∼ age`.

Despite its triviality, toy example I imparts three important lessons. First, not conditioning on a confounder may result in biased or even spurious estimation, and conditioning on a child of the explanatory variable may lead to compromised precision. Second, the performance of the model alone cannot be used to judge its adequacy: we could easily be misled by the robust statistical evidence of the tree-girl association if we overlook and fail to condition on the confounding influence of age (Fig. 6A). On the other hand, we could also be misled by the strong statistical evidence of the girl growth rate in the model (2), `girl ∼ tree + age`, when we simply condition on the tree height (Fig. 6B). Third, it is important to align the model with the effect of interest: distinct effects may warrant the adoption of separate models. The model (2), `girl ∼ tree + age`, is adequate for assessing the tree-girl association in height, but it is a poor candidate for estimating the girl growth rate.

#### 3.2.2 Revisiting toy example II

The relationship between height and weight in Q2 of toy example II also illustrates the consequences of not conditioning on a confounder. Previously, it was uncertain regarding which estimation among the three models in Fig. 3 (also shown in 7B) was more accurate. However, as depicted in the DAG in Fig. 7A, sex is a confounder, so it should be included as a covariate. If it is not conditioned on, the estimation from the simple regression (7), `weight ∼ height`, will be biased. We will later discuss the subtle differences between the other two models that include sex as a covariate, one with and the other without the interaction.

**Figure 7.**
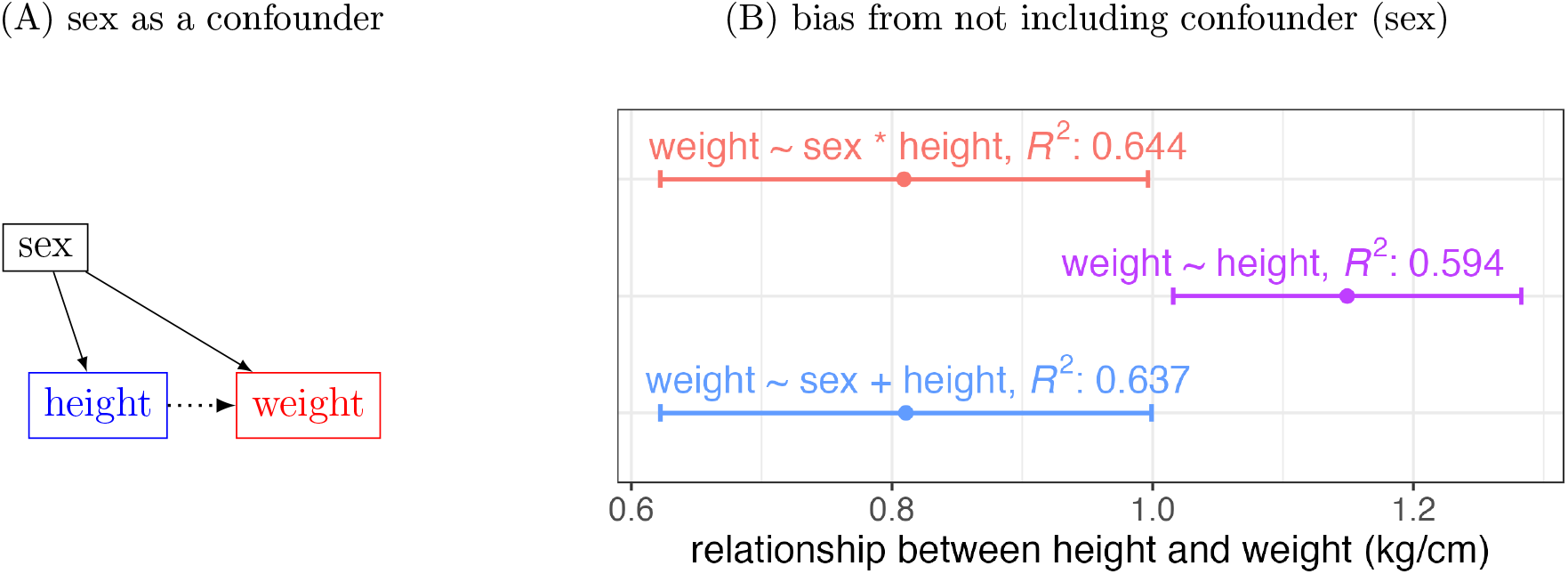
Revisiting the association between height and weight in toy example II. (A) The DAG portrays sex as a confounder between height and weight. The dotted arrow highlights the relationship between the explanatory variable of height (blue) and the response variable of weight (red), with each solid arrow indicating the a priori influence direction. (B) Fig. 3C is repeated here, demonstrating the bias introduced by failing to condition on the confounder, sex (purple).

### 3.3 Variable type: collider

It is essential to avoid conditioning on a collider. From the DAG perspective, a path with a collider is noncausal and closed (*X* → *C* ← *Y*, Fig. 4C). Conditioning on collider *C* effectively adds the noncausal path between *X* and *Y*, leading to biased estimation. *Selection bias* is actually one common manifestation of inappropriately including a collider, and this may produce a counterintuitive association between *X* and *Y*, also known as Berkson’s paradox (Pearl and Mackenzie, 2018). In general, biases may occur in various directions such as overestimation, underestimation, masking a true effect, sign flipping, and spurious effect (Appendix A.2).

#### 3.3.1 Toy example III: academic publications and selection bias

Consider a scenario where the decision to accept or reject manuscripts hinges on two primary aspects: the trending nature of the research topic, and the level of quality and rigor in the study. We then pose the question: *How does the trendiness relate to the level of rigor in research manuscripts?* Suppose that we gather data from thousands of peer-reviewed publications within the literature. It is quite likely that we would observe a negative correlation between these two variables (McElreath, 2020; Fig. 8B). However, does this correlation necessarily imply that researchers choosing popular topics tend to compromise on thoroughness, while those investigating less trendy subjects are inherently more meticulous?

**Figure 8.**
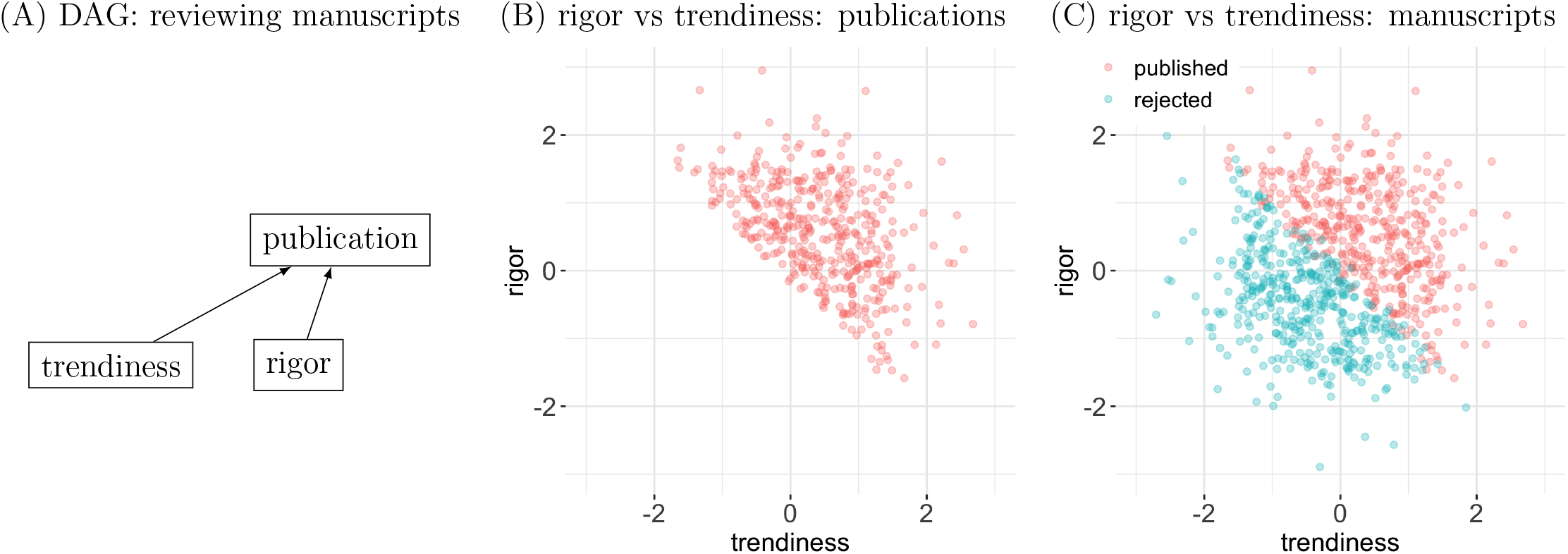
Schematic illustration of collider bias in academic publications (McElreath, 2020). (A) Suppose that during the reviewing process, the decision of accepting/rejecting manuscripts is based on the combination of two scores: rigor and trendiness. The relationships among trendiness, rigor, and publication are shown in the DAG. The reviewing process plays the role of a collider. (B) The consequence of the selection bias during the reviewing process is the negative association between rigor and trendiness among publications, as schematically illustrated among 400 publications. (C) When all manuscripts are presented, no obvious association exist between rigor and trendiness.

In fact, a negative correlation due to selection bias often arises when the data is restricted to the combination of two selected, but not necessarily correlated, features (e.g., rigor and trendiness). For academic manuscripts, the variable marking the acceptance decision is a collider between trendiness and rigor, as shown in the DAG in Fig. 8A. If we condition on the acceptance of manuscripts (Fig. 8B), one would get the impression that the rigor of manuscripts is negatively associated with their trendiness. This is precisely the scenario described above, where we conditioned on published manuscripts to characterize the relationship. On the other hand, with all manuscripts considered, little association would likely show (Fig. 8C), and the negative correlation in Fig. 8B due to selection bias would be considered spurious.

We emphasize that the presence of colliders often affect the accuracy, as well as the generalizability, of results; simply increasing sample sizes will not remove their effects and or decrease the likelihood of misinterpretation. Within neuroimaging, generalizability based on consortium data gathered from prospective studies (e.g., UK Biobank, ABCD) will still be susceptible to collider bias, such as from uneven representativeness and attritions (e.g., missing data) across different groups (Munafò et al., 2018; Cheetham et al., 2023; Schoeler et al., 2023; Gard et al., 2023). It is important to note that studies based on these large datasets can certainly *avoid* collider bias, but merely that explicit care should be taken when filtering subjects and dealing with large numbers of potential covariates.

#### 3.3.2 Revisiting toy example II

In Q3 of toy example II, we examined sex differences in height with three possible models, each of which dealt differently with including weight as a possible covariate. We can draw a DAG for these variables to provide a principled perspective. Since weight is influenced by both sex and height, it is a collider (Fig. 9A), and should *not* be conditioned on. Fig. 9B shows the estimates for the sex difference of height in the various scenarios. These findings aptly showcase the biases and compromised precision when a collider is included as a covariate.

**Figure 9.**
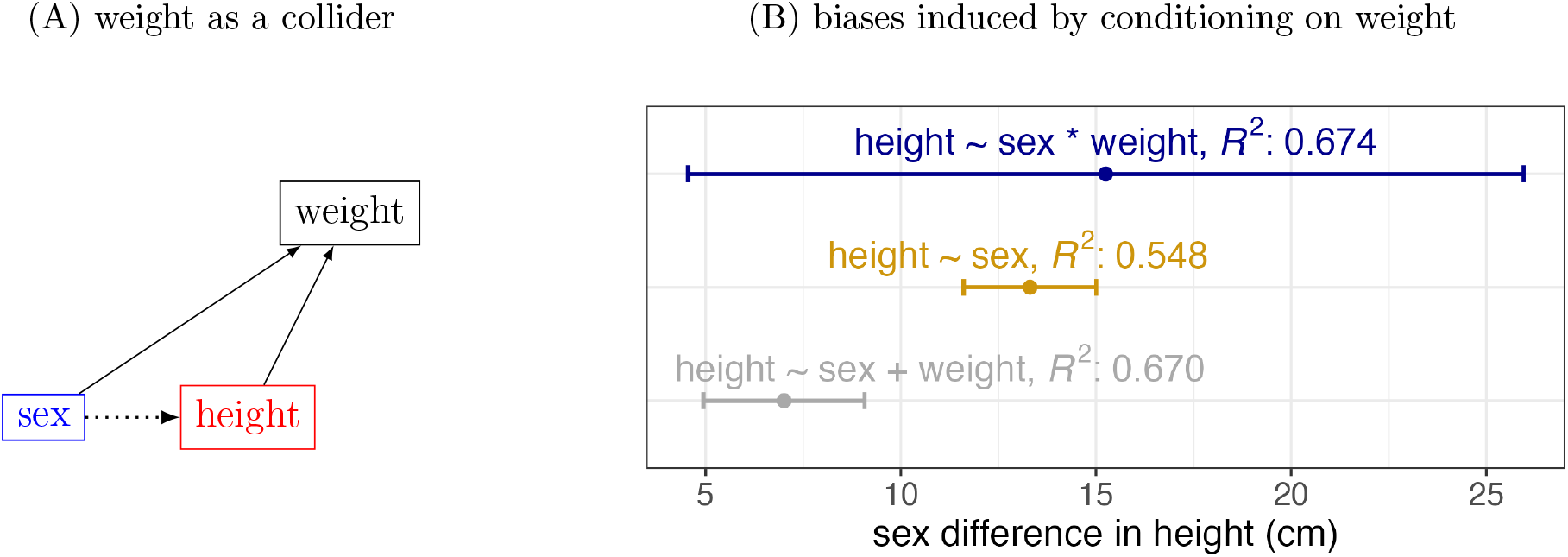
Revisiting the sex difference in height in toy example II. (A) The DAG portrays weight as a collider between sex and height. The dotted arrow highlights the relationship between the explanatory variable of sex (blue) and the response variable of height (red), with each solid arrow indicating the a priori influence direction. (B) Fig. 3D is repeated here, demonstrating the biases introduced by conditioning on the collider, weight.

### 3.4 Variable type: mediator

The impact of a mediator is subtle, affecting result interpretability, and should be treated with extreme caution. When a mediator is present, two causal paths exist that allow information flow from *X* to *Y* (Fig. 4B): the direct causal path *X* → *Y*, and an indirect causal path *X* → *C* → *Y*. One may be interested in the *total* effect, which contains the contribution of both causal paths, or in the *direct* effect, which contains only the direct causal path and excludes the indirect path. If the *total* effect of *X* on *Y* is desired, the covariate *C* should *not* be conditioned on; otherwise, the indirect causal path would be closed, and part of the causal effect would be missing from the estimation. On the other hand, if the direct effect of *X* on *Y* is desired, one should close off the indirect causal path by conditioning on *C*; otherwise, information would be contaminated by the indirect causal path. In general, biased estimation of the total effect from conditioning on a mediator may occur in various directions such as overestimation, underestimation, masking a true effect, sign reversal, and spurious effect (Appendix A.3).

#### 3.4.1 Revisiting toy example II

Q1 in toy example II investigates the weight difference between sexes, serving as a prototypical case of a mediator. Figure 10A illustrates the DAG that encapsulates our prior knowledge, indicating that sex influences height when considering the impact of sex on weight. This scenario naturally gives rise to two distinct types of inquiries. Firstly, one can explore the total effect: *What is the weight disparity between sexes?* Secondly, one could address a counterfactual question by focusing on just the direct effect: *Assuming we know an individual’s height, what would be the weight difference if that individual is male as opposed to female?*

**Figure 10.**
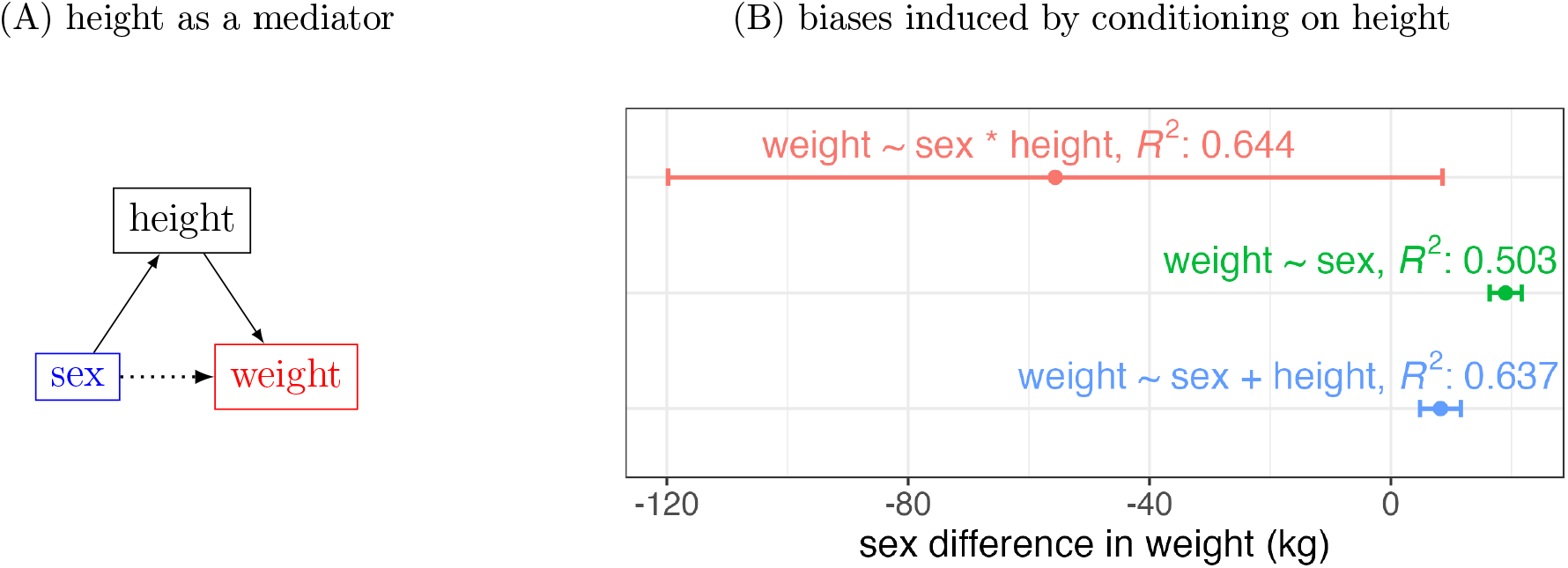
Revisiting the sex difference in weight in toy example II. (A) The DAG portrays height as a mediator between sex and height. The dotted arrow highlights the relationship between the explanatory variable of sex (blue) and the response variable of weight (red), with each solid arrow indicating the a priori influence direction. (B) Fig. 3B is repeated here, demonstrating the biases introduced by conditioning on the mediator, height.

In the presence of a mediator, researchers often seek to understand the total effect. For instance, when comparing weight between sexes, a straightforward approach is to construct a basic model of a two-sample *t*-test: `weight ∼ sex`. In contrast, introducing height as a covariate would lead to biased estimation (Fig. 10B). In the DAG language, this action effectively closes off the indirect causal path (`sex → height → weight`, Fig. 10A), adversely affecting what we would view as the total effect.

There are times when researchers may focus on the direct effect in the presence of a mediator. For example, one might be interested in understanding the influence of sex on weight when both sexes share the same height. This scenario is often framed as a counterfactual inference: given the prior knowledge of an individual’s height, what additional insight does knowing their sex provide? Estimating such a direct effect typically may require employing an analytical technique known as centering, a topic we will elaborate in the next section.

## 4 DAG applications

The preceding discussion provided a basic background for causal inference. This commentary does not aim to comprehensively cover all aspects. Extensive discussions can be found in the literature (Pearl, 2009; Elwert and Winship, 2014; Morgan and Winship, 2014; Rohrer, 2018; Cinelli et al., 2022; Wysocki et al., 2022). In typical analyses, DAGs involve numerous variables with intricate paths that are more complex than what are represented in Figures 4 and 5. In other words, constructing a comprehensive DAG that captures all potentially relevant causal relationships could easily result in an overwhelming number of variables, including latent ones.

In such cases, one may have to be realistic about the model complexity and manageability, while still avoiding modeling pitfalls.

DAGs play a pivotal role in characterizing various variable types, extending their importance beyond observational studies. Although causal inferences have traditionally found their place in fields like epidemiology, econometrics, sociology, psychology and biology, this approach has progressively infiltrated diverse domains. Focusing on causal relationships facilitates not only the formulation of accurate models but also the evaluation of model results. Below, we apply the concept of DAGs to a few modeling scenarios, including the motivating example in the Introduction, and demonstrate the importance of contextual knowledge in understanding the interrelationships among the variables and in model building.

### 4.1 Impact of reversing explanatory variable and response variable

The position and the labeling of variables in a model reflect the analyst’s prior knowledge about their relationships. This is directly relevant to the first item in the motivational example list in Sec. 1.1: What is the consequence of switching an explanatory variable with the response variable in a model? Consider a simple relationship with only two variables, in which an explanatory variable *X* is assumed to influence the response variable *Y* : *X* → *Y*. If linear, this relationship can be expressed as a simple regression:

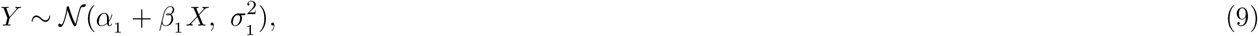

where α_1_, *β*_1_, and *σ*_1_ are the model intercept, slope, and standard deviation, respectively.

For comparison, we construct another model by reversing the two variables, which introduces a different set of parameters α_2_, *β*_2_, and *σ*_2_ :

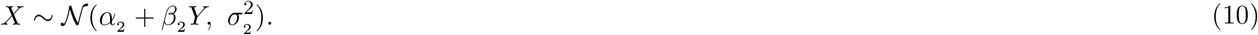

When analyzing a dataset, we have the option to adopt either model and estimate the corresponding parameters. It can be argued that the sample Pearson correlation *r* between the two variables of *X* and *Y* remains the same regardless of the adopted model. Furthermore, if one solely focuses on statistical evidence, the two models provide the same *t*- or *F*-statistic values. However, even seemingly immutable correlation and statistic values may vary between the two models, particularly when covariates are introduced into the analysis. Moreover, the numerical algorithm of ordinary least squares minimization, as commonly used in conventional statistics, yields different estimates for the two slopes,

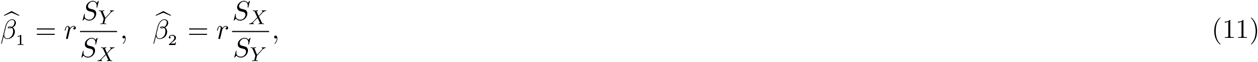

where *SX* and *SY* are the sample standard deviations of *X* and *Y*, respectively. As 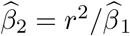, these estimates are not reciprocally related, except in a very special case: we only have equivalence 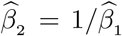 if and only if |*r*| = 1. For all other cases, the effect estimate would differ due to the opposite labeling of variables.

What causes this observed asymmetry? The dissimilarities between these two models (9) and (10) run deeper than their surface dissimilarities. By taking the algebraic perspective of their respective optimization procedures, we can unearth these distinctions. Alternatively, one can approach a regression model as a prediction challenge, where ordinary least squares plays a role as a type of regularization or penalty. Put it in another way, the biased estimation in the model (10) can be attributed to *Y* being an endogenous variable.

Here, we examine the two models from the lens of causal inference. A statistical model should be constructed based on its underlying causal assumptions, rather than solely on its output. Both models (9) and (10) can technically offer an estimate of the relationship between variables *X* and *Y*. However, neither of these models can independently justify their own output. Instead, model choice is guided by the effect of interest and the causal relationships associated with it. The first model, where we regress *Y* on *X*, implies an information flow from *X* to *Y* with an error term that represents a stochastic process, encapsulating the data variability that cannot be entirely accounted for by *X*. From the DAG perspective, this error term can be considered as a parent of *Y* (Fig. 11A). In contrast, the regression of *X* on *Y* assumes an information flow from *Y* to *X* with an error term as a parent of *X* (Fig. 11B). As a result, these two models imply distinct data generating processes, and the appropriate one should be that which most closely represents our prior knowledge. The special case in which the coefficients of *β*_1_ and *β*_2_ are symmetric occurs only when |*r*| = 1 with *no* measurement error in the response variable *Y*. In addition, the distortion in effect estimation by the second model would be exacerbated in the presence of covariates.

**Figure 11.**
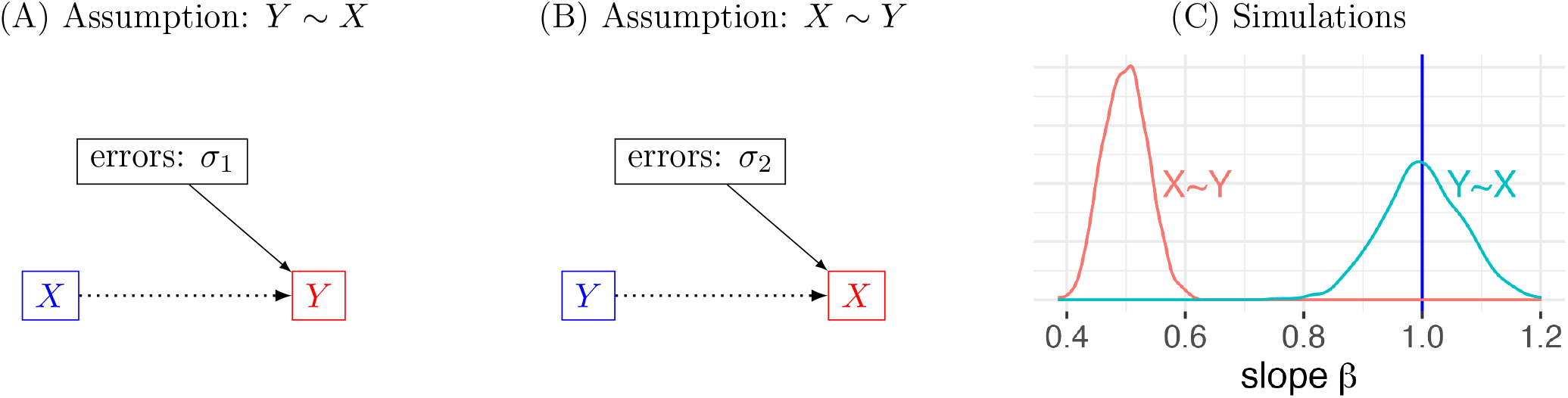
The impact of reversing an explanatory variable and a response variable. (A) The regression of *Y* on *X* assumes that *X* influences *Y* with errors representing variability in *Y* that could not be accounted for by *X*. (B) The regression of *X* on *Y* assumes that *Y* influences *X* with errors representing variability in *X* that could not be accounted for by *Y*. (C) Biased estimation may occur when a model assumes a causal relationship contrary to the reality. Under the data-generating process (9), 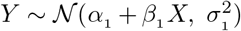 with *X* ∼ 𝒩 (0, 1), *α*_1_ = 0, *β*_1_ = 1, *σ*_1_ = 1, the adoption of the model (10), 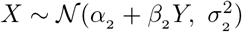, leads to substantial underestimation. Density plots are based on 1000 simulations.

Simulations prove to be a valuable tool in highlighting the distinction between the two models. As depicted in Fig. 11C, it is evident that an incorrectly specified model, characterized by swapping the explanatory variable and the response variable, may introduce a bias towards zero in effect estimation. This underestimation is closely associated with the concept of “regression dilution” or “regression attenuation,” stemming from inaccuracies in the explanatory variable (Fuller, 2006). This observation serves as a poignant reminder of a fundamental principle: a model, by itself, cannot rectify an unsuitable specification or affirm its validity. Instead, constructing an appropriate model requires the domain knowledge about variable relationships.

In Q2 of toy example II, the association between height and weight also showcases the importance of differentiating the explanatory variable and the response variable. It is reasonable to assume that height influences the weight, and not the other way around. When modeled as such, the association is estimated as 0.81 kg/cm with a 95% interval (0.62, 1.00) (Fig. 3C) based on the model (4), `weight ∼ sex + height + sex:height`. On the other hand, if we switch height and weight, the alternative model (8a), `height ∼ sex + weight + sex:weight`, would lead to an effect estimate of 0.36 cm/kg with a 95% interval (0.27, 0.44), which equates to 2.78 kg/cm and 95% interval (2.27, 3.70). This is a substantial overestimation compared to the estimate of 0.81 kg/cm with a 95% interval (0.62, 1.00) (Fig. 3C) based on the model (4).

The comparison between the two models can be framed as a sensitivity analysis. To assess the robustness of causal conclusions to changes in assumptions, model specifications, or data perturbations, sensitivity analysis evaluates the extent to which the results of a causal analysis are influenced by various factors and to identify potential sources of bias or uncertainty. In general, the analyst may modify the DAG under study, comparing the effect estimates to gauge sensitivity to alterations in underlying assumptions. This approach aids in comprehending how varying assumptions affect conclusions about causal relationships. In this specific scenario involving the causal relationships of *X* → *Y* and *Y* → *X*, where the former represents the assumed causal relationship, the model comparison illuminates the consequences of adopting an incorrect causal relationship.

### 4.2 Enhancing intepretability through covariate centering

Covariate centering is a crucial analytical technique in data analysis. Rather than directly using the values of a quantitative covariate, this method involves subtracting a specific number from the covariate values before incorporating them into the model. This number can be the mean, mode, median, or a value of particular interest to the analyst. With a rich history (Bedeian and Mossholder, 1994; Kraemer and Blasey, 2004; Gelman et al., 2020), this technique offers at least two distinct advantages. First, it enhances conceptual interpretability. Without centering, the intercept is linked to the covariate at 0, often lacking practical utility. But with centering, the intercept becomes associated with a covariate value that holds practical meaning. This convenience of interpretability extends even to categorical variables, as seen in our coding strategy for sex, with −0.5 and 0.5 assigned to females and males in models such as (4)-(6). Second, centering addresses concerns related to multicollinearity. For example, product interactions typically exhibit high correlation with their main effects. Centering also provides the added benefit of enhanced numerical stability by reducing the extent of collinearity.

#### 4.2.1 Covariate centering through the lens of causal inference

To illustrate the crucial role of centering, consider the sex difference in weight in Q1 of toy example II, where height is included as a covariate. The challenge with the model (4), weight_*i*_ ∼ 𝒩 (α + *β*_*s*_ sex_*i*_ + *β*_*h*_ height_*i*_ + *β*_*X*_ sex_*i*_ : height_*i*_, *σ*^2^), is the following. It effectively estimates the sex difference in weight at an extrapolated point where both sexes have a height of 0 cm. Consequently, the resulting estimate of −56 kg, with a 95% interval of (−120, 8) (Fig. 12A,E) lacks practical interpretability. This interpretation issue can be technically resolved through centering. Specifically, we refine the original model by introducing a centering parameter:

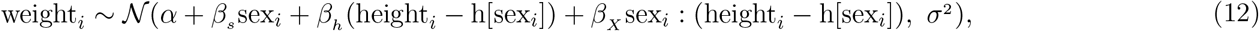

where h[sex_*i*_] is the center value for each sex. This height-centered model allows one to make two types of inference. First, when we set h[sex_*i*_] to the within-group mean, then *β*_*s*_ is interpreted as the sex difference in weight at each group’s mean height (Fig. 12B), resulting in an estimate of 19.0 kg with a 95% interval (16.7, 21.3), close to the two-sample *t*-test (Fig. 3B,E). Alternatively, if we choose a common center (e.g., the overall mean or a specific height), *β*_*s*_ represents the sex difference in weight, assuming that both sexes share the same height at this center value. This second centering method addresses a counterfactual question: when the height is known, what extra information could sex provide about weight difference? For instance, at a common height of 190 cm, the sex difference in weight is estimated as 15.2 kg with a 95% interval (7.5, 22.9) (Fig. 12E).

**Figure 12.**
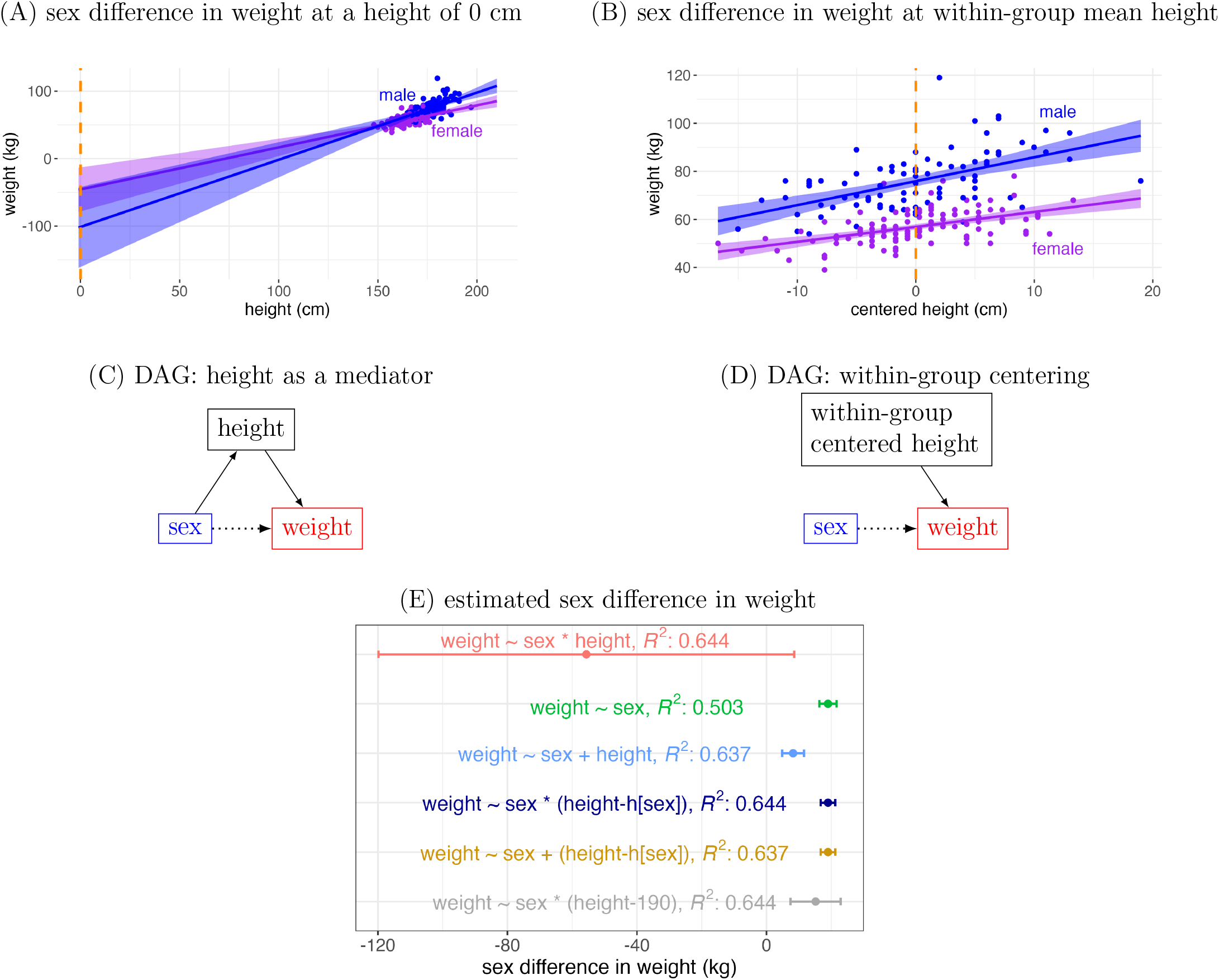
Weight difference between sexes in toy example II revisited. (A) The estimated difference between sexes based on the model (4), `weight ∼sex + height + sex:height`, is associated with the height of 0 cm (orange dashed line). (B) Centered at each sex’s mean, the two groups are shifted and aligned at their respective center (orange dashed line). The estimated effect corresponds to the sex difference in weight at each sex’s mean height. (C) Height is a mediator between sex and weight. (D) Upon within-group centering, height is no longer a mediator, but a parent of weight. (E) The estimates of sex difference in weight are compared among six models. The notation h[sex] indicates within-group mean.

Causal inference offers a fresh perspective into the centering technique. When the explanatory variable is a categorical variable, centering artificially shifts and aligns the original data to the center value. In toy example II’s Q1 with height as a mediator (Fig. 12C), one should not include height as a covariate when focusing on the total effect. However, we know a priori that males are on average taller than females (as shown in the relationship of sex → height in the DAG). Centering effectively removes the causal arrow from sex to height (and blocks the indirect causal path sex → height → weight in the DAG). Thus, this process changes the original role of height from a mediator to just a parent of the response variable (Fig. 12D). More specifically, the total effect can be obtained through *within-group centering*, because each group’s new height variable has zero mean. As height is a mediator, the direct effect can be estimated through *cross-group centering*, wherein a common center is removed from both groups. In other words, centering can convert the originally inappropriate model for total effect to one that can estimate either total the effect or direct effect. When dealing with multiple covariates, it is essential to individually justify the inclusion of each one, as well as the decision to adopt a centering strategy.

Centering can be conceptualized as a process of resolving collinearity issues. A mediator is typically correlated with the explanatory variable because the latter influences the former. Centering effectively reparameterizes the model and reduces the level of correlation. Covariate effect is often parameterized as an additive effect in common practice, without considering potential interactions and nonlinearity. In the case of the sex difference in weight in toy example II, if the interaction between sex and height is not considered, the model weight_*i*_ ∼ 𝒩 (α + *β*_*s*_ sex_*i*_ + *β*_*h*_ height_*i*_, *σ*^2^) leads to a substantial effect underestimation, with an estimated weight difference of 8.2 kg and a 95% interval (4.8, 11.6) (Fig. 12E). From the causal inference perspective, this underestimation is due to the inclusion of a mediator when the total effect is the focus. On the other hand, we consider the following model through within-group centering,

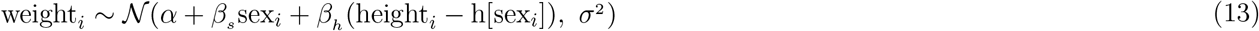

where h[sex_*i*_] is set as the average height for each sex. Estimation improvements become evident. First, centering removes the arrow from sex to height and legitimizes the inclusion of height as a covariate. Moreover, this height-centered model yields an estimate of 19.0 kg with a 95% interval (16.7, 21.3), which is comparable to the result of the two-sample *t*-test as well as that of the height-centered model with interaction (Fig. 12E). That is, while effectively reducing the correlation between sex and height, centering enhances estimation accuracy.

#### 4.2.2 Different modeling strategies

An effect of interest can typically be estimated using various models. Consider the total effect of sex difference in weight in Q1 of toy example II. We have discussed three different models: two-sample *t*-test (5), the height-centered GLM (12) with interaction, and the height-centered GLM (13) without interaction. Since uncentered height is a mediator, we do not include it as a covariate, adopting the most straightforward approach for total effect, the two-sample *t*-test. On the other hand, centering converts height from a mediator to a parent of the response variable (weight). Both GLMs, as alternative approaches, offer estimations that are similar but slightly more precise–though the improvement is negligible–when compared to the two-sample *t*-test (Fig. 12E). The difference between the two GLMs hinges on whether the interaction should be included, a subtlety we will address in a sequel of this commentary.

Model selection also occurs in estimating the association between height and weight in Q2 of toy example II. Sex is a confounder (Fig. 7A); thus, it should be included as a covariate. However, we still need to decide whether to incorporate the interaction between sex and weight. ^4^. The decision regarding the inclusion of the interaction between sex and height revolves around model parameterization. This choice hinges on the additional research interest in exploring potential differences in the slope effect of height between the two sexes. In this particular case, both approaches yield nearly identical estimations (Fig. 7B). Nevertheless, the inclusion of the interaction lends to more informative inference about the interaction: that of sex difference in the association between height and weight.

Centering may also help simplify the modeling process. Suppose that we are interested in three effects: the total effect between sex and weight, the association between height and weight, and the interaction between sex and height. For the first effect, since height is a mediator, we would normally avoid it as a covariate, adopting a simple model such as a two-sample *t*-test. For the association between height and weight, considering sex as a confounder, we would simply adopt a GLM such as `weight ∼ sex + height` or `weight ∼ sex + height + sex:height`. For the interaction, we would choose the second model. However, through centering we can merge these three separate processes into a common model (12), estimating all three effects under a unified modeling framework.

The conventional mediation analysis may help dissect the total effect into direct and indirect (mediation) effects. In toy example II, the estimated total effect for the sex difference in weight is 19.0 kg with a 95% interval (16.7, 21.3) (Fig. 12E). Mediation analysis reveals a direct effect (via the direct causal path `sex → weight`, Fig. 12C) of 7.9 kg with a 95% interval (4.2, 11.0) at the cross-group mean height of 170.6 cm and an indirect effect (via the indirect causal path `sex → height → weight`, Fig. 12C) of 10.9 kg with a 95% interval (7.97, 14.1). In essence, when both males and females share an identical height of 170.6 cm, males weigh approximately 7.9 kg more than females. This weight difference slightly varies when their height is smaller or larger (Fig. 12B). An additional 10.9 kg in the total weight difference of 19.0 kg is attributed to the mediation effect of height (i.e., males being taller than females). However, caveats exist regarding mediation analysis (Rohrer et al., 2022). For example, a confounder between the mediator and the response variable may bias the estimation of direct and indirect effects.

#### 4.2.3 Subtleties of centering

Some cautions should be practiced in centering. One subtlety pertains to the selection of center values. For toy example II, to directly compare with the two-sample *t*-test result, mean-centering can be adopted in the GLM (12). In this case, the height distributions are approximately symmetric for both sexes (Fig. 3A), making mean-centering a suitable choice. However, in other scenarios, data could be skewed, and it might be more appropriate to choose a center based on the mode, median, or a particularly meaningful value. Adaptivity in centering is crucial for accurately representing the characteristics of the data under investigation.

Centering may not always be feasible or applicable in the presence of a mediator. When the mediator has a discrete distribution (e.g., binomial) among groups, or when the explanatory variable is a quantitative variable rather than an individual-grouping factor (e.g., sex), centering is inapplicable. For instance, consider a neuroimaging study where sleep quality serves as an explanatory variable for BOLD response, and memory performance acts as a mediator. Since both sleep duration and memory measure are quantitative variables, there is no effective centering method on sleep quality that could justify the incorporation of memory performance as a covariate. In such cases, it is advisable to refrain from including a mediator if the primary interest lies in the total effect.

### 4.3 Selection biases in neuroimaging

Colliders can and do occur in neuroimaging studies. The controversial practice of “double dipping” or the circularity problem is such an example (Fig. 13A). The issue pertains to a scenario in which the same data are utilized for both selecting regions of interest or conducting analyses and later for evaluating the statistical evidence of the findings. This practice can result in inflated estimates, potentially resulting in misleading conclusions (Vul et al., 2009; Kriegeskorte et al., 2009). The neuroimaging field has largely been aware of the negative impact. Nevertheless, we can still instructively view it through the causal inference lens. Take a task-based experiment as an example, and picture the task as an explanatory variable and the full analytical results as the response variable. In the primary stage of analysis, one may select a small proportion of data based on statistical thresholding, and then proceed with a follow-up stage such as correlation analysis with behavioral data or classification. The thresholded data is a common effect between task and the full results, and plays the role of a collider. Therefore, this style of analysis contains heavy biasing for the actual relation of interest (typically overestimating the strength of relation). The earlier warnings effectively alerted practitioners of the potential risk of selection biases.

**Figure 13.**
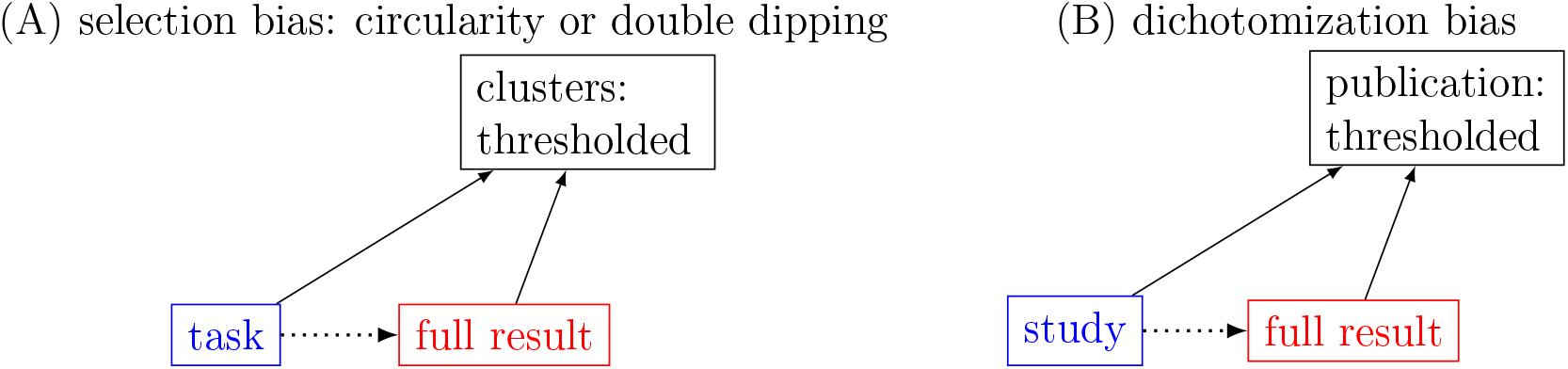
Two collider examples in neuroimaging. (A) Selection bias occurs when clusters of spatial units (e.g., voxels) are selected by thresholding the statistical evidence. (B) Publication bias may occur due to statistical thresholding in result reporting. Due to the concern about the multiple testing issue, rigorous thresholding is typically practiced. This selection bias is further deteriorated with the whole-brain results reduced to single “peak” voxels.

However, another form of selection bias often lurks within common neuroimaging practices. There has been a strong emphasis in neuroimaging on the strictness of result reporting, particularly due to concerns about the multiple testing issue. As an isomorphic counterpart of the “double dipping” case, the thresholding step plays the role of a collider (Fig. 13B). Consequently, selection bias resulting from applying stringent thresholding becomes evident from the causal inference perspective, commonly known as publication bias or the file drawer problem.

The consequences of such selective reporting methods lead to severe reproducibility challenges. For instance, subsequent meta-analyses are unavoidably biased and distorted due to the combination of three factors: selection biases due to the filtering process through stringent thresholding, the reduction of the full results to peak voxels, and the absence of effect quantification. As a result, findings that are largely consistent might likely be misperceived as differing, fueling views of a reproducibility crisis. Addressing the issue of selection bias, stemming from the emphasis on stringency and the artificial dichotomization, can be ameliorated through alternative approaches. These include placing a greater focus on model quality and effect estimation rather than decision-making solely based on statistical evidence (Chen et al., 2022), and adopting a “highlight, don’t hide” reporting strategy for studies, where more information is preserved and thresholding is de-emphasized (Taylor et al., 2023).

### 4.4 Revisiting the motivating example

We now return to the motivating example in Sec. 1.1 and try to understand through the causal inference framework. The central objective is to explore the relationship between gray matter density (GMD) and short-term memory (STM). Fig. 14 illustrates the DAG representation we conceive for the interrelationships among the seven variables. Through this DAG, the assumptions and domain expertise governing the underlying data-generating process are effectively delineated and communicated. Specifically, we postulate a causal link from GMD to STM, and from intracranial volume (ICV) to GMD. In addition, sex is known to influence ICV, GMD, and weight Ruigrok et al., 2014; Ritchie et al., 2018). Furthermore, the apolipoprotein E (APOE) genotype’s impact extends to both STM (Mondadori et al., 2007) and weight (Huebbe et al., 2015). Additionally, age is assumed to affect GMD, ICV, and STM.

**Figure 14.**
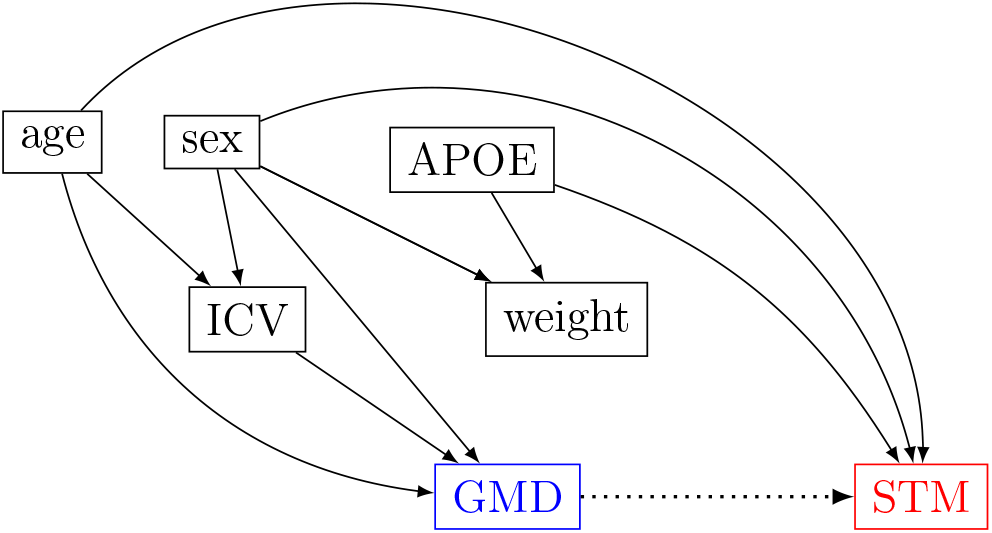
DAG for a structural MRI analysis. The research focus is on the association (dotted arrow) between gray matter density (GMD) (blue) and short-term memory (STM) (red). Five potential covariates are considered: sex, intrcranial volume (ICV), apolipoprotein E (APOE) genotype, weight and age. The relationships among the seven variables are based on subject knowledge.

We now directly address the four questions raised in the Introduction.

#### 1) Explanatory variable versus response variable

The behavioral measure STM yields a single value per individual, while the structural measure GMD is estimated at each spatial element (voxel) within a 3D image of each individual. Conventional analytical software typically operates with 3D images as the response variable, which may lead one to consider constructing a model with GMD as the explanatory variable and STM as the response variable. However, from a cognitive standpoint, it is imperative to recognize that brain structure exerts a causal influence on an individual’s behavior. Therefore, it is more conceptually sound to consider the causal direction of GMD influencing STM (GMD → STM). Conversely, estimating the causal relationship of STM influencing GMD (STM → GMD) lacks causal plausibility. As discussed in Section 4.1, a model constructed with the causal assumption of STM → GMD would result in biased estimation.

#### 2) Covariates

We expound our decision about each variable based on the DAG presented in Fig. 14.

- Age is in an open noncausal path (`GMD ← age → STM`). As a confounder, it should be included as a covariate.
- Sex, similar to age, lies in an open noncausal path (`GMD ← sex → STM`), and is also a confounder. Thus, it should be conditioned on.
- APOE only influences the response variable, but not the explanatory variable. As a parent of STM, its inclusion improves estimation precision.
- Weight is a collider for sex and APOE through a noncausal, closed path (`GMD ← sex → weight← APOE → STM`). However, given that sex acts as a confounder and should be conditioned on, incorporating weight as a covariate would neither introduce bias nor offer any benefit.
- ICV only influences GMD, but not STM; thus, it is not on any causal or noncausal path. As a parent of the explanatory variable GMD, it should not be included as a covariate.

As a result, we may adopt the following model,

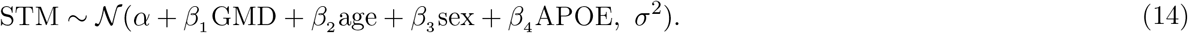

#### 3) Result interpretability/reporting

Although the model (14) can technically provide assessments for all parameters, not all estimates for sex, GMD, APOE, and age are directly interpretable. Each model is constructed within the framework of a DAG to estimate a specific relationship between the explanatory variable and the response variable. Hence, the parameter estimates for the covariates might not be appropriate due to potential bias. This is because if the focus switches to another explanatory variable or response variable, the causal relationship could undergo changes (e.g., a previous confounder might become a mediator). As a result, it may be necessary to develop a new model with a revised set of covariates to effectively capture these alterations. For instance, when estimating the total effect of sex on STM, we may condition on sex and APOE, but not on GMD (mediator). Consequently, the parameter estimate for age from the model (14) should not be reported unless the direct effect is the focus. The same issue applies to the parameter estimate of age.

#### 4) Experimental design

Since weight neither provides additional benefits nor poses any harm, omitting weight from data collection could result in potential resource savings and model simplification. On the contrary, if the investigator hypothesizes that the hours of sleep before the experiment could impact participants’ STM performance, collecting sleep data may enhance estimation precision. This is because the quality of sleep serves as a parent variable influencing the response variable.

## 5 Discussion

In this commentary, we began with a motivating scenario and two illustrative toy examples. Toy example I, which focuses on tree-girl height, demonstrated that introducing a covariate can help improve interpretability. However, it also highlights a drawback of doing so: the precision of the estimated effect for the covariate itself within the same model is poor. This suggests that one might need to employ a separate model for each effect. Toy example II of sex-height-weight offered a more intricate array of interrelationships among variables. It aided in understanding three fundamental DAG structures: confounder, collider, and mediator. Specifically, sex has the role of a confounder between height and weight, weight serves as a collider between sex and height, and height acts as a mediator between sex and weight. Importantly, not all parameter estimations derived from each model are necessarily interpretable or meaningful, further underscoring the necessity of constructing a model for each particular effect of interest.

We then applied the DAG perspective to several real-world applications. First, we showed how reversing the relationship between an explanatory variable and the response variable can lead to misestimation. Second, we underscored the role of centering, which enhances interpretability by essentially transforming a mediator into a parent of the response variable. This not only simplifies the modeling process but also enables the derivation of multiple effect estimations from a unified model. Third, we argued that the prevalent practice of strict thresholding in result reporting and preparation for meta-analyses shares the same selection bias problem as the “double dipping” issue. Lastly, we brought our discussion full circle by addressing the initial questions posed in the motivating example.

By presenting examples, terminology, and theoretical background, we have explored how causal inference offers a crucial framework for the model-building process. The examples provided in this work aim to enhance the understanding of covariate selection and uphold information integrity in result reporting more broadly. With these foundational concepts established, our future work will address neuroimaging-specific challenges in data processing and modeling, including outliers, head motion, network-level inference, and other pertinent topics.

### 5.1 DAG: the lingua franca for causal inference

In statistics, the primary objective is to construct a model that most faithfully represents the system being studied. Here, we discuss how the relationships among variables are pivotal in constructing an accurate model that mirrors the underlying data-generating process–this is the underlying raison d’être for causal inference. The model’s validity should rely on our understanding of the data generation process, specifically the causal relationships among relevant variables. This critical information, however, is not entirely encapsulated by the data itself nor inherently embedded within the model. Hence, it is imperative to seamlessly incorporate this essential knowledge as prior information during model construction. In essence, model construction should be guided by domain expertise, allowing this recognized principle to steer the covariate selection process. This approach is preferable to relying solely on statistical metrics. While a DAG provides a framework for illustrating causal relationships among variables based on current knowledge, it does not inherently preclude estimation biases due to two key reasons: 1) domain knowledge may evolve over time, prompting updates to the DAG, and effect estimation involves constructing models with specific parameterizations. However, failing to adhere to this principle carries the inherent risk of constructing a model that is fundamentally flawed, leading to potential misconstructions and misinterpretations of the underlying data-generating process. A DAG, serving as a non-parametric graphical depiction of the data-generating process, elucidates the intricate interconnections and assumptions embedded within the considered variables. Much like architectural blueprints provide simple and efficient representations for construction, DAGs serve as a standardized tool that enhances communication by systematically categorizing and classifying diverse relationships among variables, grounded in contextual knowledge. DAGs offer a structured framework for organizing pre-existing knowledge, articulating the investigator’s assumptions, and fostering comprehension for the recipient. The explicit and transparent attributes of DAGs establish a shared foundation for communication, laying the groundwork for constructive debates, critiques, and comparisons of alternative DAGs and models. This clarity and openness not only facilitate a deeper understanding but also encourage a collaborative discourse that refines and advances the overall comprehension of the underlying data structure and relationships.

With DAG representations, guidance on the inclusion or exclusion of a covariate can be simplified to a few principles. The following rules should be adopted when considering variables, in order to generally improve model quality and avoid biased estimation:

1. Confounders *should* be included.
2. Colliders *should not* be included.
3. Mediators have an extra consideration, depending on the nature of the effect of interest:
  a. Mediators *should* be included for direct effect.
  b. Mediators *should not* be included for total effect.
4. Children/descendants of the response *should not* be included.
5. Children/descendants of the explanatory variable, when included, may hurt estimation precision.
6. Parents/ancestors of the response, when included, may hurt estimation precision.
7. Parents/ancestors of the explanatory variable, when included, improves estimation precision.

Finally, we list some common varieties of biased estimation that can arise in covariate selection:

- *Confounding bias:* a confounder is omitted from the analysis.
- *Mediator bias:* a mediator is omitted even though a direct effect is of interest.
- *Spurious mediation bias:* a mediator is included even though a total effect is of interest.
- *Collider bias:* a collider is included.

It is noteworthy that biases stemming from the first two scenarios align with the concept of “omitted variable bias” in econometrics (Clarke, 2005).

The DAG approach shares both historical and methodological connections with path analysis and structural equation modeling. Path analysis aims to estimate direct and indirect relationships between variables within a causal model (Kline, 2023), achieved by explicitly parameterizing the model through path coefficients, assuming linear relationships, and normally distributed residuals. However, with both assumptions and model specifications represented in a single process, its utility is confined to linear models, resulting in a simpler and more limited scope that encounters challenges with complex causal structures (Rohrer et al., 2022). In contrast, DAGs graphically represent causal relationships, offering a visual framework for understanding mechanisms. They explicitly articulate causal assumptions and enable analysts to identify confounders, mediators, and colliders. Unlike path analysis, DAGs are considered nonparametric because they do not prescribe model specification, parameterizations, or distributional assumptions. Analysts may select a model that accurately reflects causal relationships, incorporating nonlinearity and a diverse range of distributions (e.g., binomial, Poisson, or log-normal). While path analysis remains an option if deemed suitable, DAGs offer a more flexible and comprehensive approach.

### 5.2 Implications of causal inference

Covariate selection should not rely solely on statistical measures. Traditionally, covariates have been evaluated based on their “importance,” quantified by metrics like *p*-values, amount of variance, or correlation with other variables. While it might be tempting to simplify and automate the decision process using a single measure, this approach essentially prioritizes statistical indicators over fundamental causal relationships, akin to allowing the “tail to wag the dog.” In essence, covariate selection should be rooted in the principles of causal inference.

Causal inference based on DAGs holds significant implications not just for model construction but also for experimental design. Using causal inference, the model can be established before any data are collected, and indeed can inform the data selection process. Traditionally, experimental design in neuroimaging encompasses aspects like jittering, randomization, inter-trial intervals, and trial and participant sample sizes. As shown in the motivating example, employing causal thinking through DAGs brings to the forefront potentially involved covariates simultaneously. Furthermore, taking this proactive step before data collection would prevent situations where data for crucial observable variables (e.g., confounders) are not collected, while avoiding the unnecessary allocation of resources to covariates that may not be needed (e.g., colliders).

Causal inference is also pivotal for result interpretation and reporting. The research question influences the role of each variable: a variable may be simultaneously a mediator, a collider or a confounder depending on the effect of interest, and it can play a different role in separate research questions using the same data. These considerations will dictate different analytical strategies, but again these will be established and guided by causal inference principles. In other words, each model centers on a specific relationship between an explanatory variable and the response variable. When a covariate becomes an explanatory variable, an alternative model might be necessary. For instance, the parameter estimate for a confounder may not be causally interpretable due to two primary reasons. First, the explanatory variable (e.g., height in Fig. 4A) acts as a mediator when the confounder (e.g., sex in Fig. 4A) becomes an explanatory variable. Second, confounders of the confounder may be absent in the current model. Consequently, the results obtained from one particular model might not always be inherently interpretable or meaningful; knowing this prevents a researcher from accidentally making claims that have no causal basis. Apart from the primary effect of interest, researchers might feel compelled to present all other parameter estimates along with the associated statistical evidence in a separate table. However, this practice is problematic and has led to the coining of the term “Table 2 Fallacy” (Westreich and Greenland, 2013; Keele et al., 2020; Hünermund and Louw, 2023).

Causal relationships confer greater interpretive power than simple association analysis. While association analysis focuses on identifying relationships between variables, it does not establish causation nor provide a clear causal direction. Despite the adage “correlation does not imply causation,” association analysis can reveal interesting patterns for further investigation and serve as a precursor to causal inference. However, the lack of a principled approach for covariate selection in association analysis leads to arbitrary and inconsistent practices, undermining the rigor and interpretability of the results. In contrast, explicitly framing association analysis within the causal inference framework ensures that covariate selection is guided by causal principles, leading to a more rigorous and interpretable approach (Hernán, 2018). Integrating causal inference principles into association analysis can enhance the overall quality of research by improving rigor and interpretation power, aligning with the idea that addressing causation-related concerns strengthens the research process.

The pivotal role of statistics in scientific investigations necessitates careful reflection. In practice, statistics significantly shapes crucial processes like model construction, result interpretation, and reporting. A delicate balance often emerges between the rigor of statistical models and the insights derived from domain expertise.

However, the undue dominance of statistical models, often perceived as quantitative and objective gatekeepers, tends to diminish the importance of domain knowledge. Covariate selection serves as a pertinent example, where conventional practices prioritize a covariate’s accountability of data variability or its correlations with the explanatory and response variables. Similarly, strict thresholding in common practice allows modeling outcomes to unilaterally dictate reporting decisions. Consequently, a blind reliance on statistical evidence can inadvertently overshadow valuable contextual knowledge, leading to substantial biases.

Our demonstration here highlights the necessity for a statistical model to align closely with the underlying data-generating process. Validation of a model’s validity and the credibility of its results should not be independent but rooted in an understanding of the intricate web of causal relationships and underlying mechanisms. Contrary to a binary perspective of “statistical significance,” statistical evidence exists on a continuous spectrum, challenging the common practice of rendering modeling outcomes definitive based solely on arbitrary thresholds. A more informed approach involves considering the continuum of statistical evidence alongside domain-specific knowledge, which includes prior research findings and anatomical structures, to reach robust conclusions.

It is essential to recognize that domain knowledge and statistical models are not antagonistic forces but rather should be viewed as complementary components. Models should not be perceived as independent arbiters relying solely on raw data, where terms like “data-driven” might erroneously equate to reliability, even though data can inherently carry biases. Results generated by models should not be unquestionably accepted, especially when they contradict established prior information. Instead, there should be a collaborative synergy between domain knowledge and statistical models. Contextual insights ought to guide model selection and validate outcomes, while statistical models refine and update existing knowledge. Importantly, a DAG serves to clearly articulate model assumptions and specifications. Including DAGs in publications would significantly facilitate comparisons across studies, ignite intriguing discussions, and contribute broadly to advancing domain understanding.

### 5.3 Consequences of overlooking causal relationships

The importance of clearly stating the relationships among variables cannot be overstated. Without a causal understanding of the relevant variables, both the investigator and the reader may struggle to properly interpret the subtleties of a model, as well as possible discrepancies between models that use different covariates. The following issues are prevalent in common practice:

#### 1) Lack of justification

Analysts are usually aware of the importance of conditioning on confounders. However, the awareness of categorization and distinctions among the various types of relationships among variables remains largely lacking. Due to conceptual confusions, covariates’ impacts are often simply treated as generically confounding effects without considering causal relationships. Various factors like brain size, physiological fluctuations (e.g., breathing rate, heart rate), age, scanning sites, scanners, and software packages are lumped together and termed “confounding effects” without proper context and justifications. This oversight highlights a broader lack of awareness regarding the nuanced interplay among variables.

#### 2) Using model indices as justifications

The term “causal salad” describes thoughtlessly including diverse covariates in a model without thorough consideration (McElreath, 2020). This approach purports causal relationships to variables based on their inclusion and strong statistical evidence. Consequently, there is a prevalent tendency to interpret changes in estimates and statistical metrics (e.g., p-value, *R*^2^, information criteria) as indications of causation.

#### 3) Covariate mishandling

When variables of no interest are simply added as covariates, it is especially problematic when they act as mediators, collilders, or descendants of the explanatory variable. In the presence of a mediator, *mediation analysis* (Lee et al., 2021) *has been developed through a complex set of steps to parse various effect components. To avoid pitfalls, it is preferable to avoid conditioning on a mediator unless a clear justification for direct–instead of total–effect can be presented*.

Many analysis malpractices fall into this bias estimation pitfall. When intuition is largely absent or obscured, misinterpretation due to the inclusion of mediators can easily occur. For instance, consider a study where different groups (e.g., children and adults) and conditions (e.g., congruent and incongruent) exhibit differences in reaction time. Similarly, covariates such as brain volume, head size, cortex surface area, thickness (Ruigrok et al., 2014; Ritchie et al., 2018), weight, height, and body-mass index show differences between sexes. Consequently, simply adding these variables into a model, as commonly practiced, can lead to biased estimation. When not properly justified, the availability of various covariates in large datasets (like ABCD and UK Biobank) further exacerbates the likelihood and severity of this malpractice.

The potential pitfalls of mishandling mediators apply to both within-individual and between-individual covariates. It is generally easier to recognize the subtleties of a between-individual covariate acting as a mediator (e.g., head size in the context of the relationship between sex and BOLD response). However, similar nuances also come into play with within-individual covariates. Take the example of two conditions of easy and difficult tasks that are associated with short and long durations (or reaction times) at both the trial and condition levels. In this scenario, caution is warranted when dealing with duration (or reaction time) as a covariate. If the primary focus is on contrasting the two conditions (i.e., total effect), one should either omit the covariate from the model or employ appropriate centering strategies (e.g., within-condition centering) when incorporating the covariate. Adding the covariate in the model without proper centering would address a counterfactual and likely unintended question: What is the BOLD response difference when easy and difficult tasks have the same duration (or reaction time)? These subtleties substantially affect analyses at both the individual and population levels, even if they often go unnoticed in the field.

#### 4) Result reporting

Parameter estimates from a model are frequently reported without adequate justification. In a model involving more than one explanatory variable, it is possible to simultaneously derive estimation for all parameters. Consequently, various estimates are often reported from a single model that encompasses multiple covariates. As illustrated here, each effect of interest is associated with a specific model, whose design considerations are tailored to the specific pair of explanatory and response variables, potentially making the estimation for another parameter estimated from the same model unsuitable. In simpler terms, different effects may necessitate distinct models.

#### 5) Artificial dichotomization

In data analysis, a pervasive practice involves subjecting results to stringent thresholding. This artificial dichotomization introduces a substantial reproducibility challenge. The widespread insistence on strict thresholding in result reporting within neuroimaging stems from concerns about multiple testing. Paradoxically, this dichotomization approach shares a common bias issue with the well-acknowledged problems associated with “double dipping”. That is, dichotomization contributes to a reproducibility crisis, leading to distorted meta-analyses influenced by publication bias. Discarding sub-threshold voxels before meta-analysis is akin to excluding low-significance individuals before conducting group analysis–a practice that can be likened to estimating an iceberg based solely on its tip. While researchers are typically wary of the latter practice due to its potential to skew results, the same caution should be exercised in the former case.

The issue of reproducibility is poignantly illustrated by looking at the NARPS project. About 70 teams analyzed the same dataset, and when strict thresholding was applied before comparisons, conclusions were judged to be inconsistent across a large fraction of teams (Botvinik-Nezer et al., 2020). Even though this conclusion of inconsistency has been most widely cited and broadcast in literature and social media, we can recognize that thresholding actually plays the role of a collider, and so conditioning on stringent thresholding produces a strong selection bias, strongly distorting interpretation. In fact, without inserting dichotomization, the very same results were evaluated as being predominantly consistent (Taylor et al., 2023; Botvinik-Nezer et al., 2020). Moreover, instead of solely reporting statistical values through artificial dichotomization, greater information can be conveyed by emphasizing the quantification of effect magnitude and its associated uncertainty, as demonstrated in the two toy examples here (e.g., Figs 1-3, 6).

Some may contend that biases are an inevitable trade-off essential for effectively controlling false positives. However, this price is disproportionately inflated, primarily due to the oversight of anatomical hierarchies within the brain in the conventional mass univariate approach (Chen et al., 2020; Chen et al., 2022). Consequently, the overemphasis on controlling for false positives leads to excessively high false negatives. In addition, given that studies with results falling short of excessively stringent thresholding remain concealed from the literature, the level of bias can be insidiously high. Nevertheless, addressing this issue is possible by reporting effect estimates (e.g., percent signal change, Chen et al., 2017) and embracing a slightly modified approach — adopting a “highlight, don’t hide” strategy. This entails maintaining a certain level of rigor through soft thresholding while visually presenting the remaining data with some degradation (Taylor et al., 2023; Chen et al., 2022). This nuanced approach not only acknowledges the need for evidence stringency but also mitigates the potential distortion introduced by hiding full results. Relying solely on single studies for certainty is rarely effective. Promoting information integrity and transparency, rather than black-and-white thinking through stringent gatekeeping, is key to fostering reproducibility.

### 5.4 Suggestions

Causal inference demands a departure from conventional modeling methodologies. The adoption of Directed Acyclic Graphs (DAGs) proves invaluable in organizing existing knowledge and establishing a rationale for the inclusion of covariates. This approach encourages researchers to meticulously contemplate the interconnections among variables, visually articulate the reasoning behind their model selection, and contribute to the ethos of open science and research reproducibility. Here, we summarize our recommendations for the modeling process:

#### 1) Clear research hypotheses

Explicitly define research questions and effects of interest. Identify explanatory and response variables for each effect, listing all relevant observable or latent variables. Consider potential constraints on data collection.

#### 2) DAG presentation

Integrate DAGs into both experimental design and model construction. Apply causal thinking to experimental design to optimize data collection, avoiding unnecessary resource allocation. Include latent variables in DAGs for better interpretation and future guidance. Recognize that the absence of an arrow signifies a strong assumption of no meaningful causal relationship between variables.

#### 3) Variable selection

Justify adopted models by presenting DAGs based on domain knowledge and theory. Follow principles of conditioning on confounders, avoiding conditioning on colliders. Exercise caution with mediators, explicitly choosing between investigating total or direct effects. Adopt conventional mediation analysis when comparing total and direct effects (VanderWeele, 2016). Only condition on parents of the response variable among parent and child variables.

#### 4) Modeling techniques

Construct models consistent with DAG representations for each effect. Properly center quantitative variables, especially when estimating total effects in the presence of a mediator. Consider interactions and nonlinearity (Chen et al., 2021) when relevant.

#### 5) Result interpretability

Not all parameter estimates from a model are causally interpretable. Acknowledge that a single model might be insufficient for a given dataset. Guard against the “Table 2 Fallacy” by reporting only results suitable for the model.

#### 6) Transparent reporting

Enhance meta-analysis accuracy and reproducibility by avoiding strict thresholding. Report results containing both estimate magnitude and uncertainty.

### 5.5 Limitations

In typical data analysis, the intricacies of the actual data-generating process often elude complete understanding. Consequently, this process must be inferred through contextual knowledge, relevant theory, and plausible assumptions. The accuracy and precision of effect estimation hinge on the alignment of the proposed DAG with the true data-generating process. Creating and sharing a DAG serves to make underlying assumptions explicit, opening them to public scrutiny. While DAG assumptions may vary in active research domains, a consensus on key features may evolve over time. However, several limitations are associated with DAG representations:

#### 1) Dependence on domain knowledge

Research domains such as neuroimaging are inherently complex, involving numerous covariates and intricate influence patterns. A DAG serves as an abstracted representation at the research focus level, relying on the quality of prior information for its creation. The accuracy of the DAG is paramount for a causal interpretation, and while a model derived from the DAG does not guarantee precise estimation of a causal effect, it explicitly represents existing knowledge and underlying assumptions. Incorporating prior knowledge into the modeling process with a DAG helps mitigate biases, facilitating the attainment of reasonably accurate estimates with adequate precision.

#### 2) Qualitative nature

DAGs solely represent hypothesized causal relationships between variables, remaining qualitative in nature. While they aid in conceptualizing interrelationships, they do not specify how these relationships should be parameterized. The specific functional forms (linear or nonlinear), the presence or absence of interactions, and distribution assumptions must be informed by additional sources of knowledge.

#### 3) Difficulty in estimating direct effects

Estimating direct effects becomes challenging when a confounder between a mediator and the response variable is influenced by the explanatory variable. In such cases, alternative approaches must be adopted (van Zwieten et al., 2022). Conversely, when an unobservable confounder exists, the estimation for the direct effect becomes biased (McElreath, 2020). This complexity underscores the need for careful consideration in estimating direct effects.

Causal inference has flourished as a field for several decades. While our discussion here focuses primarily on covariate selection and result reporting, numerous other methodologies have emerged in recent years. These include instrumental variable analysis, inverse probability weighting, matching, and more. Readers interested in these advanced methods can explore the relevant literature, including works by Grosz et al. (2023) and Greifer and Stuart (2023), for comprehensive coverage.

## 6 Conclusions

This commentary underscores the paramount importance of adopting a causal thinking approach in statistical modeling. While statistical techniques can unveil the strength of associations between variables, they fall short of inherently revealing the underlying causal relationships. Therefore, grounding the model-building process in an understanding of causal relationships, guided by domain knowledge, emerges as a central imperative.

DAGs emerge as potent tools for intuitively and efficiently communicate complex relationships in data analysis. Through illustrative examples and simulations, we offer a primary introduction to DAGs. Recognizing the pivotal role of categorizing covariates in variable selection, we argue that a firm grasp of causal inference profoundly influences the critical stages of statistical modeling: designing experiments, constructing models based on these relationships, interpreting results, and reporting outcomes. Causal thinking aids researchers in designing experiments more efficiently, discerning underlying assumptions, and formulating theoretical hypotheses, thereby fostering analytical rigor and reproducibility. The explicit presentation of DAGs facilitates judicious covariate selection, promoting open science and transparency, allowing for external scrutiny. Readers gain awareness of the model’s inherent assumptions, empowering them to assess how these assumptions may impact the study’s conclusions.

## Acknowledgments

The research and writing of the article were supported by the NIMH Intramural Research Program (GC and PAT, ZICMH002888) of NIH / HHS, USA, andFonds de recherche du Québec – Santé Postdoctoral Fellowship (ZC).

## Appendices

### A Bias estimation arising from improper inclusion or exclusion of Covariates

Decisions regarding the inclusion or exclusion of covariates play a crucial role in statistical modeling. Three fundamental rules guide these decisions: include confounders, exclude colliders, and include or exclude mediators based on the focus on direct or total effects. In the following subsections, we illustrate the consequences of not adhering to these rules through numerical simulations.

#### A.1 Biased estimation when a confounder is not conditioned

The importance of conditioning on confounders cannot be overemphasized. Failing to do so can introduce bias, i.e., either systematic underestimation or an overestimation of the true effect within the model. A common misconception is that the absence of statistical evidence for an effect eliminates concerns about confounding factors. While scenarios that produce overestimation may be more familiar (Fig. 7B) and spurious association (Fig. 6C), it is also possible that the failure to condition on a confounder can lead to potential underestimation, concealment of a genuine relationship, or even a change in its direction (Fig. 15). This underscores the subtle and challenging nature of situations where confounders are either unobservable or unrecognized.

**Figure 15.**
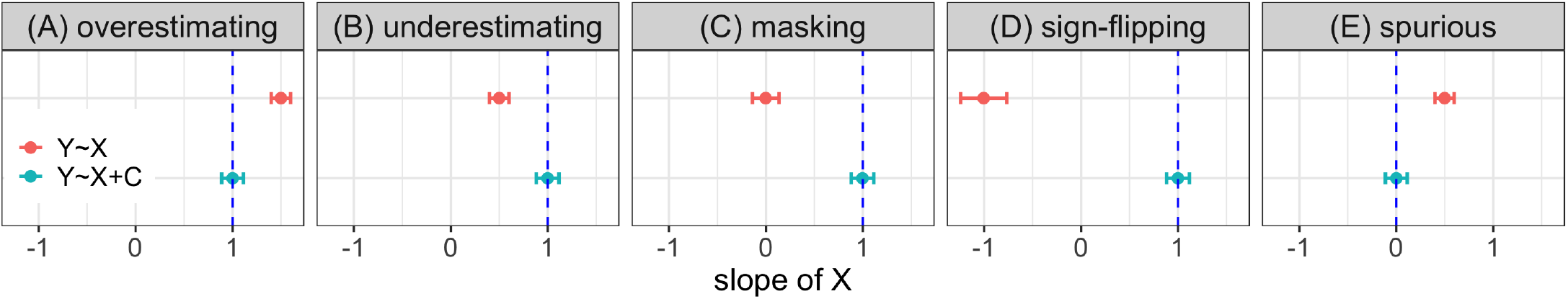
A confounder, when not conditioned, can bias estimation in various ways. Consider the scenario where *C* acts as a confounder between the explanatory variable *X* and the response *Y* with the data-generating process: *C* 𝒩 (0, 1), *X*∼𝒩 (*β*_0_ *C*, 1), *Y*∼ (*β*_1_ *X* + *β*_2_ *C*, 1). Simulations for the slope of *X* indicate the following five scenarios relative to the ground truth (blue vertical dashed line) when the confounder *C* is not conditioned on (red): (A) overestimation (*β*_0_ = 1, *β*_1_ = 1, *β*_2_ = 1); (B) underestimation (*β*_0_ = −1, *β*_1_ = 1, *β*_2_ = 1); (C) masking the presence of the association (*β*_0_ = −1, *β*_1_ = 1, *β*_2_ = 2); (D) changing the sign of the association (*β*_0_ = 1, *β*_1_ = 1, *β*_2_ = 4); and (E) spurious effect (*β*_0_ = −1, *β*_1_ = 0, *β*_2_ = 1). Estimation with mean (dot) and uncertainty (error bar with two standard deviations) are based on 1000 simulations.

### A.2 Biased estimation with conditioning on a collider

The influence of conditioning on a collider can be subtle yet impactful. Berkson’s paradox (Pearl and Mackenzie, 2018) provides an insightful example. In Fig. 16, we employ numerical simulations to illustrate a range of bias scenarios: overestimation, underestimation, effect concealment, sign change, and spurious effects.

**Figure 16.**
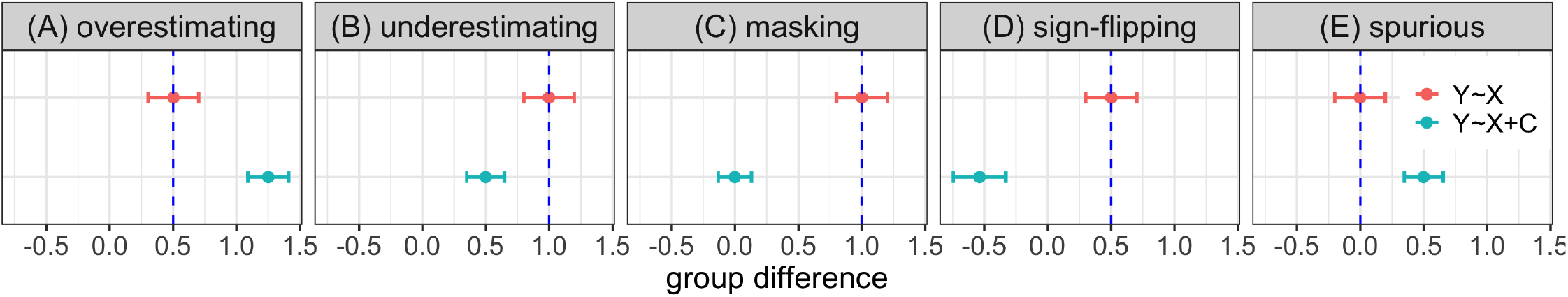
A collider, when conditioned, can bias estimation in various ways. Consider the scenario where *C* acts as a collider between a factor of group with two levels and the response *Y* with the data-generating process: *Y*_*g*_∼𝒩 (*d*_*g*_, 1), *C*_*g*_ ∼𝒩(*a*_*g*_ + *b*_*g*_ *Y*_*g*_, 1), *g* = 1, 2. Simulations for the group difference *Y*_2_−*Y*_1_ indicate the following five scenarios relative to the true effect (blue vertical dashed line) when the confounder *C* is not conditioned on (red): (A) overestimation (*d*_1_ = 0, *d*_2_ = 0.5, *a*_1_ = 2, *a*_2_ = 0, *b*_1_ = 1, *b*_2_ = 1); (B) underestimation (*d*_1_ = 0, *d*_2_ = 1, *a*_1_ = 0, *a*_2_ = 0, *b*_1_ = 1, *b*_2_ = 1); (C) masking the presence of the effect (*d*_1_ = 0, *d*_2_ = 1, *a*_1_ = 0, *a*_2_ = 0, *b*_1_ = 1, *b*_2_ = 3); (D) changing the effect sign (*d*_1_ = 0, *d*_2_ = 0.5, *a*_1_ = 2, *a*_2_ = 0, *b*_1_ = −1, *b*_2_ = −0.5); and (E) spurious effect (*d*_1_ = 0, *d*_2_ = 0, *a*_1_ = 0, *a*_2_ = − 1, *b*_1_ = 1, *b*_2_ = 1). Estimation with mean (dot) and uncertainty (error bar with two standard deviations) are based on 1000 simulations.

### A.3 Impact of conditioning on a mediator

The challenge in handling a mediator often arises from the need to distinguish between total and direct effects. Doing so improperly within models leads to distorted estimations, and doing so inconsistently across models can cause confusion or contradictions in the literature. Two common mistakes are typically encountered in practice. The first involves including a mediator as a covariate when the investigator aims to obtain the total effect. The second centers around misinterpreting an estimation as a total effect when conditioning on a mediator, leading to distorted estimation. The estimation distortion of total effect resulting from conditioning on a mediator can induce biases. Such biases happen in various ways, including overestimation, underestimation, masking real effect, sign flipping, and spurious difference (Fig. 17).

**Figure 17.**
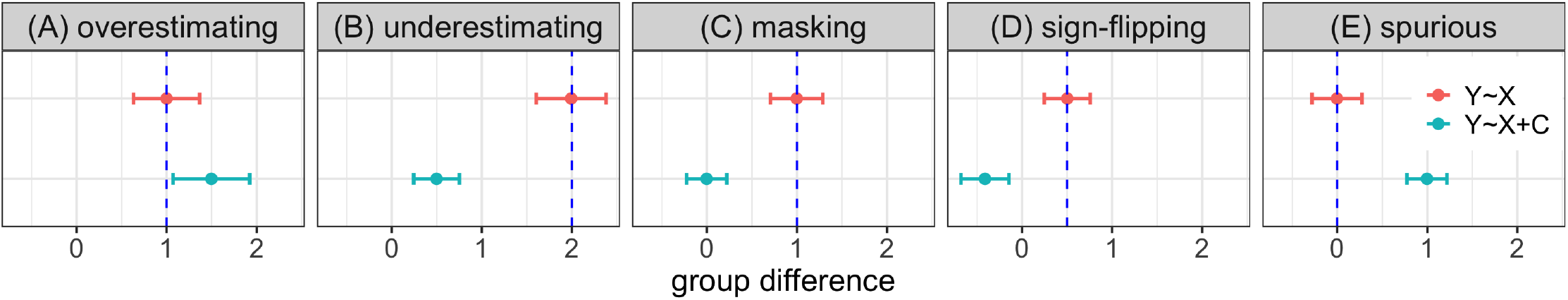
A mediator, when conditioned, can bias estimation for the total effect in various ways. Consider the scenario where *C* acts as a mediator between a factor of group with two levels and the response *Y* with the data-generating process: *C*_*g*_∼𝒩 (*d*_*g*_, 1), *Y*_*g*_∼𝒩 (*a*_*g*_ + *b*_*g*_ *C*, 1), *g* = 1, 2. Simulations for the group difference *Y*_2_ − *Y*_1_ indicate the following five scenarios relative to the true effect (blue vertical dashed line) when the confounder *C* is not conditioned on (red): (A) overestimation (*d*_1_ = 0, *d*_2_ = 1, *a*_1_ = 0, *a*_2_ = 0, *b*_1_ = 2, *b*_2_ = 1); (B) underestimation (*d*_1_ = 0, *d*_2_ = 1, *a*_1_ = 0, *a*_2_ = 0, *b*_1_ = 1, *b*_2_ = 2); (C) masking the presence of the effect (*d*_1_ = 0, *d*_2_ = 1, *a*_1_ = 0, *a*_2_ = 0, *b*_1_ = 1, *b*_2_ = 1);(D) changing the effect sign (*d*_1_ = 0, *d*_2_ = 1.333, *a*_1_ = 0, *a*_2_ = 0, *b*_1_ = 1, *b*_2_ = 0.375); and (E) spurious total effect (*d*_1_ = 0, *d*_2_ = −1, *a*_1_ = 0, *a*_2_ = 1, *b*_1_ = 1, *b*_2_ = 1). Estimation with mean (dot) and uncertainty (error bar with two standard deviations) are based on 1000 simulations.

## Footnotes

1 https://github.com/afni/apaper_DAG1

2 The pattern of estimating tree growth rate closely mirrors that of girl growth rate. Two models akin to (2) and (3) can be formulated. However, introducing girl height as a covariate compromises the precision of the estimation.

3 For the sake of intuitive and concise presentation, we adopt the Wilkinson notation for model formulas (Wilkinson and Rogers, 1973), which is also widely used in R. Specifically, the compact notation *A ∗ B* indicates that both the main effects of *A* and *B* and their interaction *A* : *B* are present in the model, i.e.: *A ∗ B* = *A* + *B* + *A* : *B*.

4 The interaction between sex and height can be conceptualized within the framework of the conventional moderation analysis where sex acts as a moderator between height and weight. Alternatively, this interaction can be seen as an additional cause influencing the response variable in the DAG (Attia et al., 2022).

